# ByeTAC: Bypassing an E3 Ligase for Targeted Protein Degradation

**DOI:** 10.1101/2024.01.20.576376

**Authors:** Eslam M. H. Ali, Cody A. Loy, Darci J. Trader

## Abstract

Targeted protein degradation utilizing a bifunctional molecule to initiate ubiquitination and subsequent degradation by the 26S proteasome has been shown to be a powerful therapeutic intervention. Many bifunctional molecules, including covalent and non-covalent ligands to proteins of interest, have been developed. The traditional target protein degradation methodology targets the protein of interest in both healthy and diseased cell populations, and a therapeutic window is obtained based on the overexpression of the targeted protein. We report here a series of bifunctional degraders that do not rely on interacting with an E3 ligase, but rather a 26S proteasome subunit, which we have named ByeTACs: Bypassing E3 Targeting Chimeras. Rpn-13 is a non-essential ubiquitin receptor for the 26S proteasome. Cells under significant stress or require significant ubiquitin-dependent degradation of proteins for survival, incorporate Rpn-13 in the 26S to increase protein degradation rates. The targeted protein degraders reported here are bifunctional molecules that include a ligand to Rpn-13 and BRD4, the protein of interest we wish to degrade. We synthesized a suite of degraders with varying PEG chain lengths and showed that bifunctional molecules that incorporate a Rpn-13 binder (TCL1) and a BRD4 binder (JQ1) with a PEG linker of 3 or 4 units are the most effective to induce BRD4 degradation. We also demonstrate that our new targeted protein degraders are dependent upon proteasome activity and Rpn-13 expression levels. This establishes a new mechanism of action for our ByeTACs that can be employed for the targeted degradation of a wide variety of protein substrates.

## Introduction

Protein degradation is an essential cellular function by which proteins are naturally destroyed in order to maintain protein homeostasis.(*1*) The rate of protein degradation differs depending on the cell type and life cycle stage.(*2*) In eukaryotic cells, the majority of protein clearance is handled through the ubiquitin proteasome system (UPS).(*3–6*) Proteasomes, which are integral components of the UPS, are made up of two main regions: the 19S regulatory particle (19S RP) and 20S core particle (20S CP).(*7, 8*) Ubiquitinated proteins are recognized by the 19S RP via three subunits: Rpn-1, Rpn-10, and Rpn-13.(*9*) The 19S RP then recruits deubiquitinates, unwinding the protein to reduce its tertiary structure, and shuttling the protein into the catalytic core (20S CP) to be degraded. Proteolysis Targeted Chimeras (PROTACs ®) were developed to harness the activity of the UPS to degrade a protein of interest. Targeted degraders offer several advantages over the conventional small-molecule inhibitors. For instance, degraders exert rapid, robust, and potentially prolonged degradation of the targeted proteins and evolution of resistance can be less prominent as compared to small molecule inhibitors.(*10, 11*) Due to their promising therapeutic advantages, research in this field has experienced significant acceleration, leading to a staggering 25+ ubiquitin-dependent degraders in clinical trials.(*12, 13*)

To date, the identification of E3 ligases capable of effectively inducing targeted protein degradation has been limited to cereblon (CRBN) and cullin ligase Von–Hippel Lindau (VHL).(*14*) This has restricted the types of proteins that can be ubiquitinated and degraded via the UPS. Also, ubiquitin-dependent degraders can lead to destruction of protein in healthy cells, which can have negative side-effects. A degradation mechanism that can be applied broadly to more proteins of interest and can obtain a better therapeutic window is of great interest.

A new approach that can induce targeted protein degradation by directly binding to a 26S proteasome subunit would alleviate the requirement to interact with an E3 ligase. This methodology could allow for a more universal degradation strategy that could potentially be applied to proteins that lack ubiquitination sites or do not interact with the E3 ligases for which ligands have been developed. A recent report demonstrated this concept by inducing a binding interaction with a 26S proteasome subunit to degrade BRD4.(*14*) Here, they developed a macrocyclic ligand for PSMD2 (Rpn1) tethered to a BRD4 ligand and demonstrated that binding to this subunit allowed for efficient degradation of their protein of interest (POI) around 1 μM. Although this proof of concept established the ability to degrade a POI in a ubiquitin-independent manner, the system is not ideal as it requires a large macrocyclic peptide binder and cannot differentiate degrading BRD4 in healthy cells versus diseased. Direct engagement of a 26S subunit that is only essential in diseased or inflamed cells could allow for selective degradation of a POI, which is lacking in current degradation techniques.

Rpn-13 (ADRM1), one of the proteins within the 19S RP that can recognize ubiquitinated proteins, is observed to be overexpressed in many cancer types and is indicated to be essential for their viability.(*15–18*) Knock-down or -out of Rpn-13 in yeast and healthy mammalian cells has no effect on viability.(*19–21*) For these reasons, Rpn-13 has emerged as a promising therapeutic target for the treatment of various cancer types including multiple myeloma and ovarian cancer.(*20, 22–24*) Several covalent binders, including RA190 and XL5, were found to interact with Rpn-13 at Cys-88, leading to tumor growth inhibition in xenograft mouse models of multiple myeloma and ovarian cancer.(*20, 25*) KDT11, a non-covalent inhibitor, was discovered as a reversible and highly selective binder of Rpn-13 that shows selective toxicity to multiple myeloma cells.(*26*)

Here we describe the development of non-covalent bifunctional molecules that bind Rpn-13 and BRD4, leading to the degradation of BRD4 though direct association with the 26S proteasome. We generated a series of degraders that recruit our POI, BRD4, with binding moiety JQ1 attached to a recently reported Rpn-13 ligand, TCL1, with varying PEG linker lengths.(*27*) Excitingly, we are able to use nanomolar concentrations of our ByeTACs for degradation of our POI through this ubiquitin-independent process. Our degraders were also shown to be most effective in cells that are highly dependent on Rpn-13 for survival. This new mechanism of targeted degradation is reliant on Rpn-13, which is required for the survival of many cancer types and can generate a significant therapeutic window. Our unique method of protein degradation can compete with current degraders at nanomolar concentrations and offers an advantage of bypassing a required interaction with an E3 ligase.

## Results

Through ubiquitin interacting domains, ubiquitinated proteins initially interact with Rpn-10 and then Rpn-13.(*28, 29*) The additional binding interactions of a ubiquitinated protein via Rpn-13 is believed to increase the rate at which ubiquitinated proteins can be degraded.(*14, 30*) Rpn-13 has been reported to be non-essential in healthy cells, but seems to enhance the proliferation of various cancer cell types.(*21, 23, 31–35*) Rpn-13 is comprised of a DEUBAD domain, a Pru domain, and an unstructured region that connects them.(*24*) The ligands to Rpn-13 previously described interact with the Pru domain of Rpn-13. We envisioned developing bifunctional molecules that could associate a non-ubiquitinated protein with the 26S proteasome through an interaction with Rpn-13, which can lead to degradation, **Figure 1A**.

**Figure 1.**
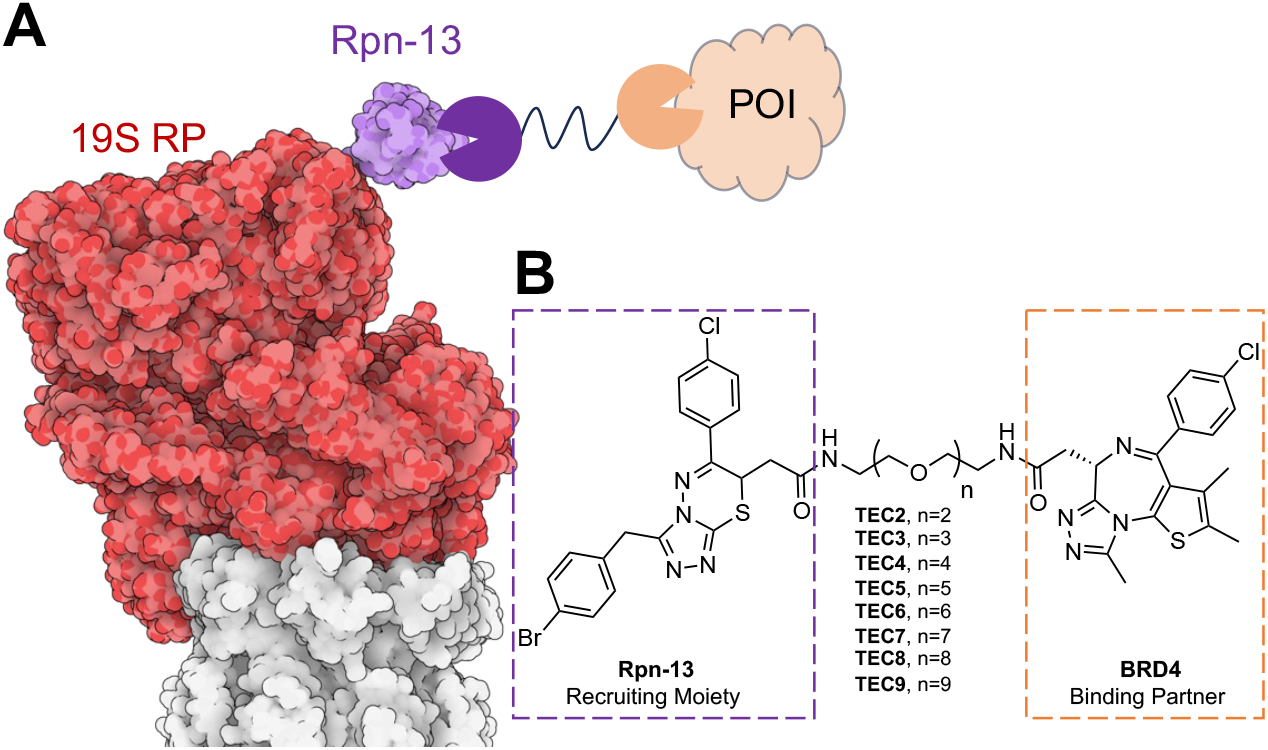
**(A)** Modified PDB structure 4CR2 to show 19S RP (red) with Rpn-13 (purple). Proposed ubiquitin-independent tethering of a protein of interest (orange) to Rpn-13 for subsequent degradation by 26S proteasome. **B)** General structure of TEC degraders with Rpn-13 recruiting moiety TCL-1 (purple) appended to BRD4 binding partner JQ1 (orange) with varying PEG linker lengths.

We recently reported the discovery of TCL1, a non-covalent binder to Rpn-13’s Pru domain.(*27*) TCL1 is shown to bind to Rpn-13 with modest affinity (K_D_ ∼ 26 μM). From this previous study, we knew that we could modify the carboxylic acid of TCL1 into an ester and still retain binding to Rpn-13. Therefore, we chose that position of TCL1 to incorporate PEG linkers with variable length appended to a moiety to interact with the POI for degradation, **Figure 1B**. In this study, we selected BRD4 as the POI. BRD4 is widely used as a model POI for the development of bifunctional degraders such as ARV-825, A-1874, dBET1, and MZ-1.(*29, 36, 37*) From these degraders we chose ARV-825, a cereblon-based BRD4 degrader, as our control to compare our ubiquitin-independent degraders.(*6*)

To generate our suite of degraders we needed to first establish a synthesis of TCL-1 from commercially available sources. Starting with 4-(4-chlorophenyl)-4-oxobutanoic acid and 2-(4-bromophenyl) acetic acid we developed an 8-step synthetic method to efficiently generate TCL-1 in enough quantity for further derivatization. To the carboxylic acid we appended commercially available PEG linkers to vary the space between the two ligands. This is important as the linker length of other degraders has been shown to be important for inducing the proper confirmation for degradation.(*38–40*) To the other end of the linker, we installed a BRD4 ligand, JQ1, generating our first series of ByeTACs (TEC 2-9).

This suite of degraders was assessed in cell lines reported to be reliant on Rpn-13 for survival in comparison to cell lines that are not as dependent on Rpn-13-mediated degradation. Ramos B-cell lymphoma cells was selected as they have been previously reported to have high Rpn-13 levels for survival, **Figure 2A, S1**, and **S2**.(*20, 26, 27*) All degraders, including ARV-825 as our control, were dosed at low nanomolar concentrations for 18 hrs before harvesting, and subjecting the cell lysate to western blotting for BRD4, Rpn-13, and tubulin (gel loading control). Rpn-13 was included with the western blotting to ensure we were not changing its levels with our degraders.

**Figure 2.**
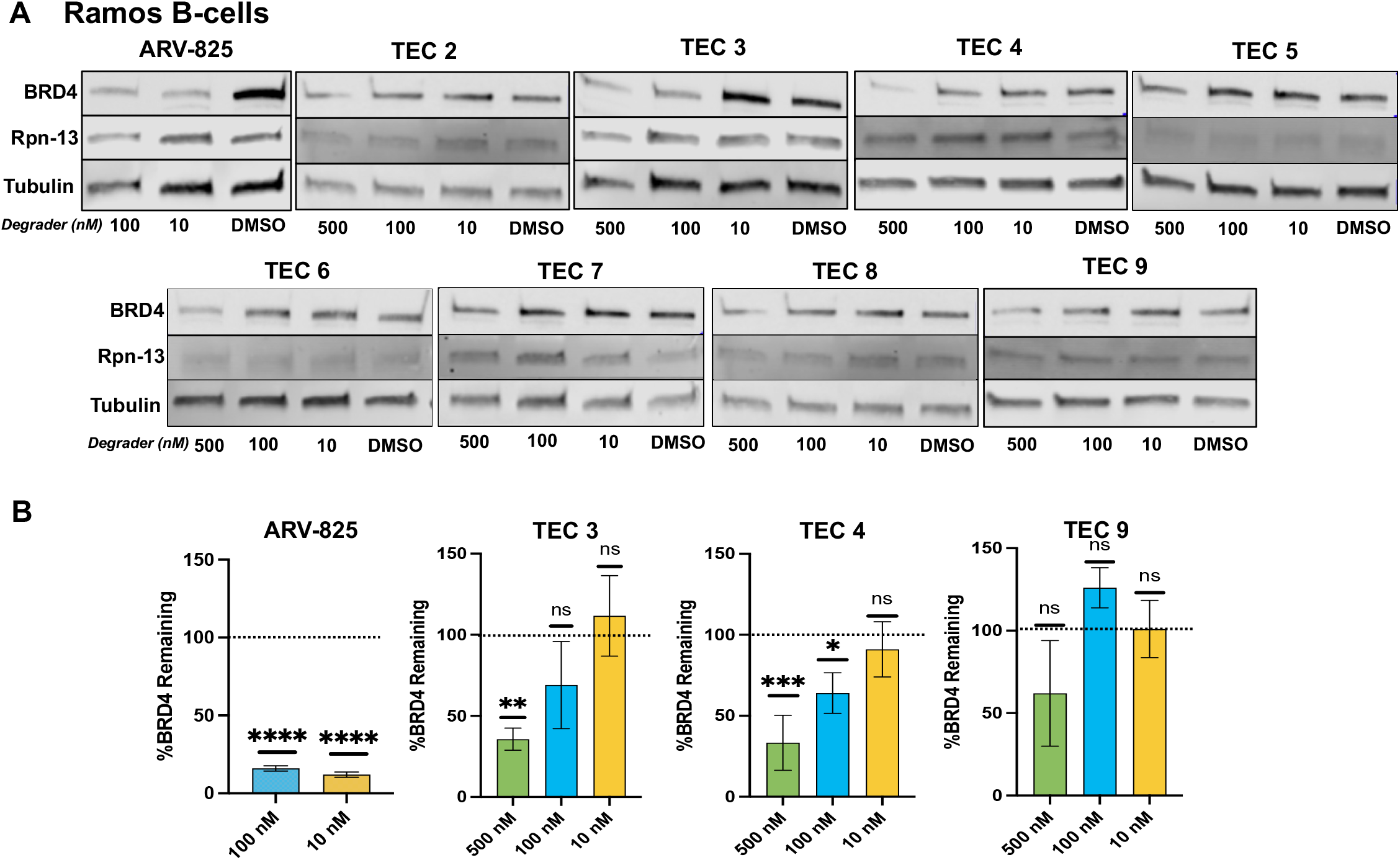
**A)** Degradation of BRD4 in Ramos B-cells with TEC degraders and ARV-825 after 18 hrs of incubation. 500 nM of ARV-825 was excluded due to its toxicity. **B)** All BRD4 bands were normalized to tubulin, and the percent BRD4 remaining calculated by dividing normalized signal by DMSO normalized signal (100%). A one-way ANOVA Dunnett’s multiple comparisons test was performed to compare each concentration to DMSO for statistical significance. p<0.0001: ****; p<0.001: ***; p<0.01: **; p<0.05: *

Excitingly, we were able to observe degradation of BRD4 in Ramos B-cells at concentrations comparable to the ARV-825 control, **Figure 2B**. We also determined the optimal linker length of n=3 (TEC3) or n=4 (TEC4) gave the most consistent and dose dependent degradation of BRD4. N=9 (TEC9) had the least amount of BRD4 degraded and would be used as our negative control for further studies. Wanting to demonstrate the versatility of this new targeted degradation method, we dosed other cell lines with TEC3, TEC4, ARV-825 as our positive control, and TEC9 as our negative control. We initially selected Raji B-cells as they have also been reported to rely on Rpn-13 recruitment for 26S survival.(*20, 27*) U87 cells were also used as this cell line has previously been reported in the development of BRD4 degraders and would be a good model for us to compare our Rpn-13 dependent method.(*41–43*) TEC3 and -4 were shown to be able to degrade BRD4 in U87 cells and Raji B-cells (**Figure 3A** and **3B**). TEC9 had the same result that was observed in the Ramos B-cells, where BRD4 degradation did not occur. Control experiments to establish that both ligands of our bifunctional degraders were required and degradation of BRD4 was proteasome dependent were conducted. No BRD4 degradation was observed if the TEC molecule was lacking one of the ligands or after inhibition of the proteasome with MG-132 (**Figure S3** and **S4**).

**Figure 3.**
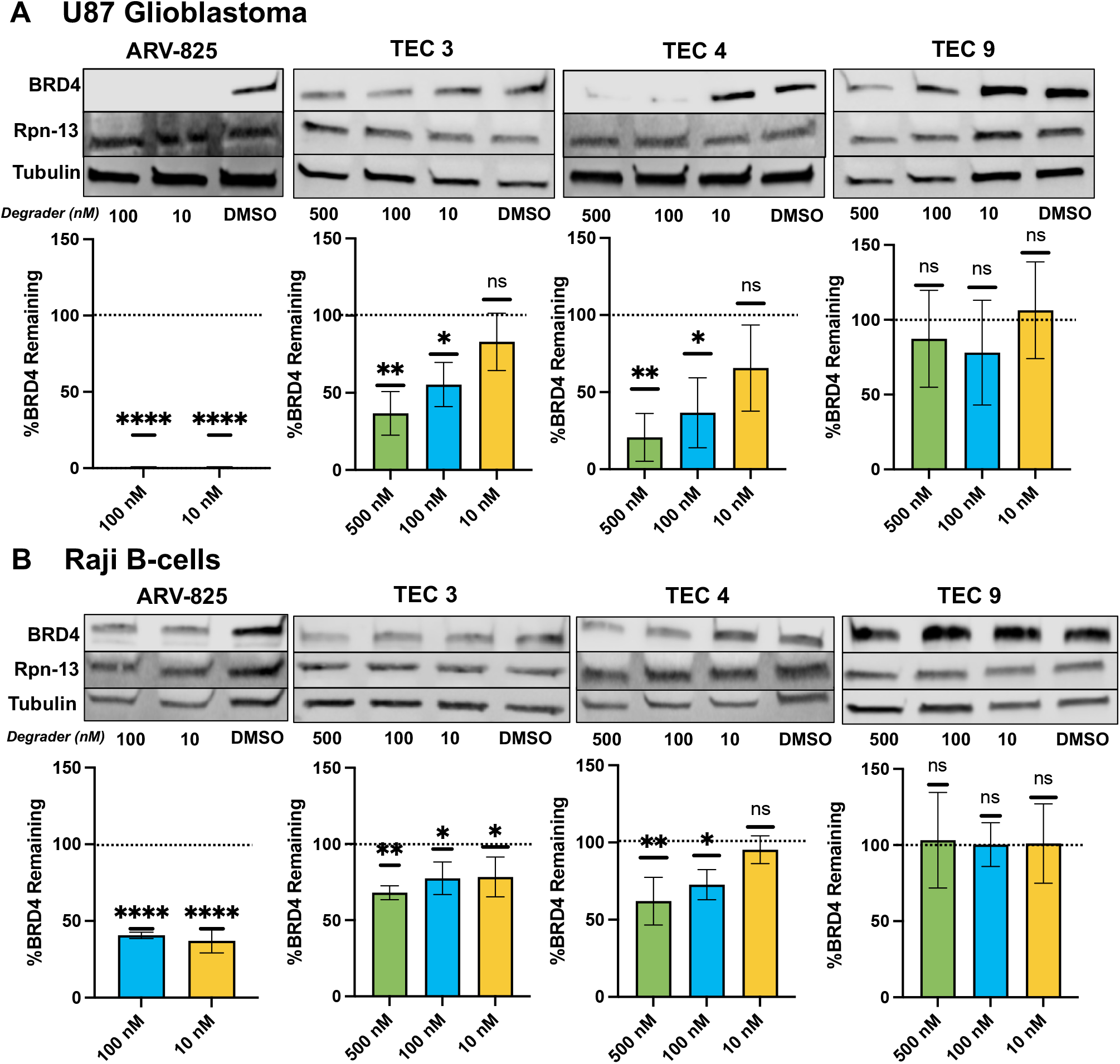
**A)** Degradation of BRD4 in U87 cells and **B)** Raji B-cells with TEC degraders and ARV-825 after 18 hrs of incubation. 500 nM of ARV-825 was excluded due to its toxicity. All BRD4 bands were normalized to tubulin, and the percent BRD4 remaining calculated by dividing normalized signal by DMSO normalized signal (100%). A one-way ANOVA Dunnett’s multiple comparisons test was performed to compare each concentration to DMSO for statistical significance. p<0.0001: ****; p<0.001: ***; p<0.01: **; p<0.05: *

HEK-293T cells were next evaluated for BRD4 degradation using our degraders. HEK-293T cells were selected as they have been reported to not rely on Rpn-13 for survival as inhibitors are not toxic.(*15, 26, 27, 44, 45*) We had hoped to observe limited BRD4 degradation which could be attributed to lower Rpn-13 expression Excitingly, we were able to see dramatically reduced to no BRD4 degradation with our best degraders, **Figure 4** and **S5**. We hypothesize that BRD4 is not degraded using our methodology as HEK-293T cells could have lower expression of Rpn-13 or 26S proteasomes in HEK-293T cells have less Rpn-13 associated with their 19S RP.(*46*) Additional studies are required to validate this hypothesis, but these results indicate we should able to obtain a significant therapeutic window between cells that are highly dependent on Rpn-13 complexed to the 26S proteasome and the healthy cell population.(*24, 45, 47*)

**Figure 4.**
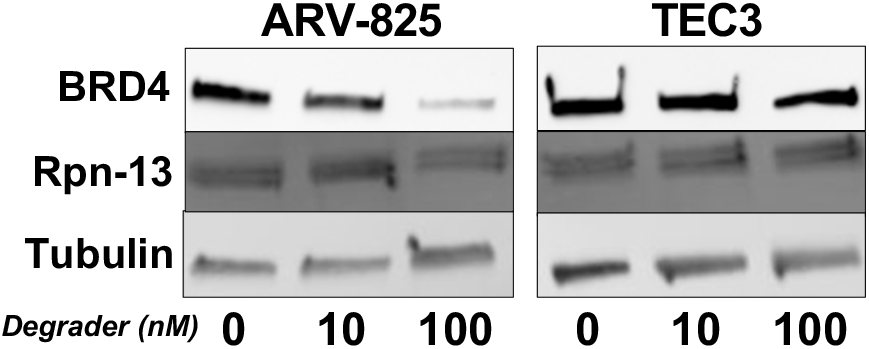
Degradation of BRD4 using ARV-825 and TEC3 in HEK-293T cells. Only ARV-825 was effective.

Thus far, we have been able to demonstrate the ByeTAC degrader series can initiate the destruction of BRD4 in cells that rely on Rpn-13 highly for survival, the degradation is proteasome dependent, and both binding moieties are required to target the POI to the 26S proteasome for degradation. However, we needed to ensure that this method of degradation is reliant on the interaction with Rpn-13. From our previous publication it is known that TCL-1 is capable of binding to Rpn-13 after modifications to its carboxylic acid,(*27*) but ensuring our full degrader is still capable of binding and facilitating degradation needed to be confirmed. A set of experiments were undertaken to confirm the reliance of Rpn-13 on the effectiveness of our degraders utilizing siRNA to Rpn-13. We expected to observe little to no degradation by our degraders after lowering the expression of Rpn-13. The ARV-825 degrader should still be able to facilitate degradation utilizing the traditional ubiquitin-dependent method and the 26S proteasome. We decided to use the U87 cells as we saw significant degradation of BRD4 at 100 nM with our degraders and ARV-825. We first optimized the conditions to knockdown Rpn-13 so that it was significant as compared to DMSO treated cells and did not lead to cell death. It was determined that with 25 nM of siRNA, we were able to significantly decrease the expression of Rpn-13 after 48 hours, **Figure S6 and S7**.

After confirmation, we dosed the U87 cells with 25 nM of Rpn-13 siRNA for 48 hours, and then dosed with either TEC4 or ARV-825 for 18 hrs at 100 nM. TEC4’s ability to degrade BRD4 was eliminated, while ARV-825 was still successful in degrading BRD4, **Figure 5**. This supports our hypothesis that the TEC degrader series require a substantial interaction with Rpn-13 to induce degradation.

**Figure 5.**
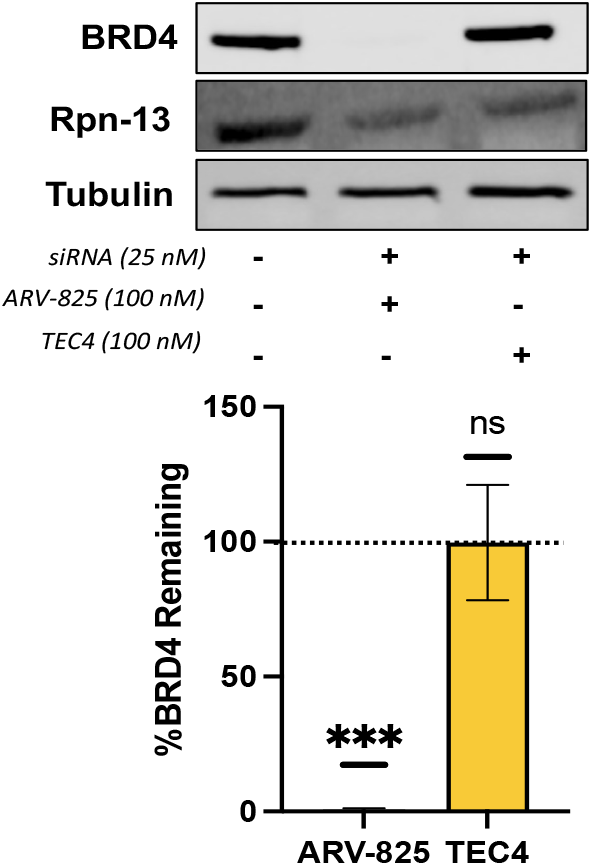
siRNA of U87 cells with ARV-825 and TEC4 at 100 nM. siRNA was dosed with 48 hrs at 25 nM, then degraders or DMSO was added for 18 hrs. A one-way ANOVA Dunnett’s multiple comparisons test was performed to compare each concentration to DMSO for statistical significance. A one-way ANOVA Dunnett’s multiple comparisons test was performed to compare each concentration to DMSO for statistical significance. p<0.0001: ****; p<0.001: ***; p<0.01: **; p<0.05: *

Upon confirmation that our method of degradation requires Rpn-13 for recruitment and subsequent degradation of BRD4, we wanted to confirm our ByeTACs were functioning through a ubiquitin-independent mechanism. We relied on the widely used TAK-243 (MLN7243), E1 inhibitor, to shut down the ubiquitin ligase cascade. TAK-243 is capable of inhibiting the ubiquitin-activating enzyme (UAE) at low nM concentrations.(*48–50*) To Ramos B-cells, TAK-243 (1 nM) was added simultaneously with ARV-825 (100 nM) or TEC 4 (100 nM). Upon treatment with the UAE inhibitor and incubation with degraders, we were able to observe TEC 4’s degradation abilities were retained while ARV-825’s was lowered, **Figure 6**. More specifically, TEC-4’s ability to degrade BRD4 when the ubiquitin ligase cascade is inhibited had only a small change (55% vs 40% when including TAK-243) while ARV-825, which requires ubiquitination of BRD4, had a diminished ability for degradation (16% versus 64%), **Figure S8 and S9**. This result helps to confirm that our ByeTAC methodology can degrade a protein without the recruitment of ubiquitin.

**Figure 6.**
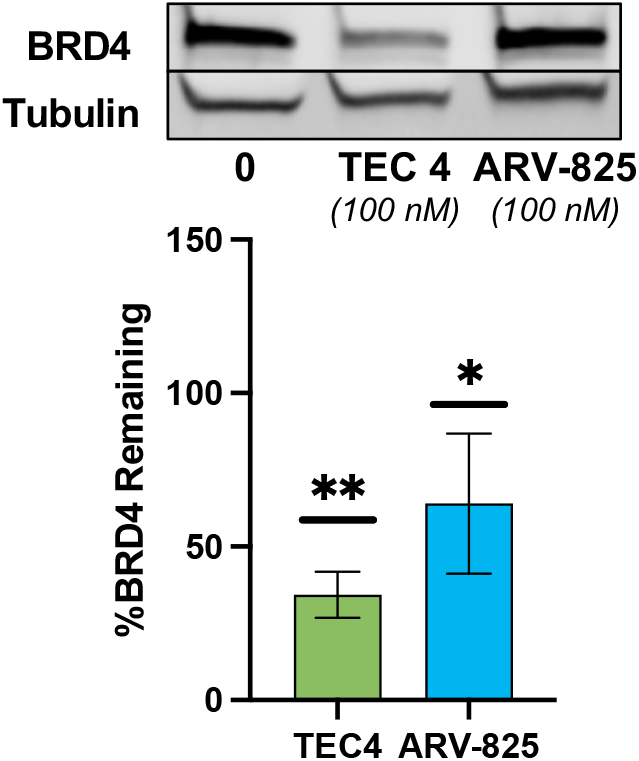
Ubiquitin inhibition of Ramos cells by TAK-243 (1 nM) followed by incubation with degrader for 18 hrs. TEC4 A one-way ANOVA Dunnett’s multiple comparisons test was performed to compare each concentration to DMSO for statistical significance. p<0.0001: ****; p<0.001: ***; p<0.01: **; p<0.05: *

Confirming that ByeTAC method of degradation is potent, dependent on Rpn-13, and can degrade a POI through a ubiquitin-independent mechanism, we wished to assess their toxicity in cell lines reported to be reliant on Rpn-13 for survival. Initially, we assessed toxicity of all eight degraders in Ramos, Raji, and U87 cells, **Figure S10**. From these cell viability experiments, we were able to see a correlation between degradation and a decrease in viability. We were able to determine that TEC3 and TEC4 were our most toxic degraders to cell lines that are reliant on Rpn-13, **Figure 7A**. We were also able to show that in HEK-293T cells, our degraders only affected viability at the highest concentration (5 μM), **Figure 7B**, which is 50-fold higher concentration than required to degrade BRD-4.

**Figure 7.**
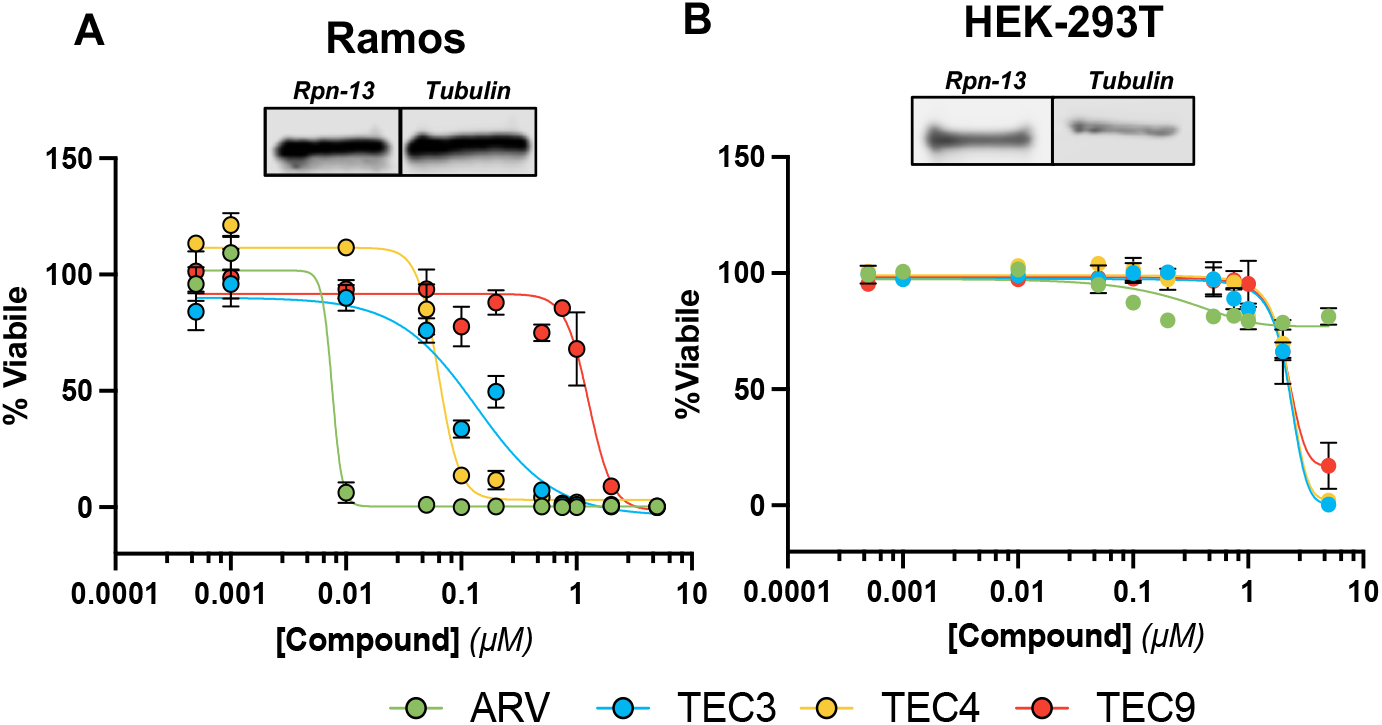
Toxicity of ARV-825 (green), TEC3 (blue), TEC4 (yellow), and TEC9 (red) in **A)** Ramos, **B)** HEK-293T cell lines. Cells were plated at 10,000 cells per well and dosed with compounds in triplicate for 48 hours. Viability was determined using CellTiterGlo® viability assay. Rpn-13 levels are also included for cells lines in singlet, triplicates are provided in supplemental material.

We also performed a number of experiments to provide evidence that our toxicity is associated with BRD-4 degradation and not from altering the parent Rpn-13 ligand, **Figure S11 and S12**.

## Conclusion

Here we have reported a new methodology for targeted protein degradation utilizing a ubiquitin-independent mechanism. This method, named ByeTACs, bypasses the needs for an E3 ligase by initiating an interaction between the POI and Rpn-13, a 26S proteasome subunit. As Rpn-13 is not essential to healthy cells, but many cancer types such as multiple myeloma, lymphomas, and ovarian are dependent upon its association with the 26S proteasome to increase the rate of ubiquitinated-protein degradation, we believe a significant therapeutic window can be obtained. This is a limited feature in traditional E3 ligase-mediated degraders. It is not just cancer cells that require high Rpn13 association with the 26S proteasome for survival, as it has been noted cells under stressed/inflammatory conditions also require it.(*34, 51, 52*) This greatly increases the disease types our ByeTACs could be effective in treating. While we have only demonstrated BRD4 degradation, we anticipate that we can exchange JQ1 for a number of other ligands. We also anticipate we can increase the potency for these degraders by incorporating different linker types and a stronger binder to Rpn-13. Overall, these studies highlight the proof of concept that degradation of a POI can be mediated through a non-covalent interaction with Rpn13.

## Associated Content

### Supporting Information

The Supporting Information is available at XXX.

### Author Information

## Acknowledgements

This work was supported by a start-up package from the UCI School of Pharmacy and the UCI Chao Family Comprehensive Cancer Center (P30CA062203). Thank you to Prof Erin E. Carlson (University of Minnesota) and Prof. Brian Paegel (University of California-Irvine) for helpful discussions.

