## Supporting Information for "ByeTAC: Bypassing an E3 Ligase for Targeted Protein Degradation"

Supporting Information Content:

**Methods**
*General Materials*
*Synthesis of TEC Degrader Series*
*General Cell Culture*
*BRD-4 Degradation Dosing Suspension Cells*
*BRD-4 Degradation Dosing Adherent Cells*
*BRD-4 Degradation Western Blot*
*Proteasome Dependency (MG-132 dose)*
*Rpn-13 (ADRM1) siRNA*
*E1 inhibition with TAK-243*
*Cellular Toxicity Assays with TEC's Suspension Cells*
*Cellular Toxicity Assays with TEC's Adherent Cells*

**Figures:**
*S1: Representative western blot for degradation studies*
*S2: Degradation of other degraders in Ramos*
*S3: Linker control degradation in Ramos*
*S4: Proteasome dependent degradation in Ramos*
*S5: Degradation with TEC4/8 in HEK*
*S6: siRNA controls*
*S7: siRNA Rpn-13 levels with degraders*
*S8 and S9: Tak-243 inhibition full gels*
*S10: Toxicity of degraders in Ramos, Raji, and U87*
*S11: Toxicity of JQ1 vs. JQ1-Linker*
*S12: Toxicity of TCL-1 vs. TCL-linker*

**Reaction Schemes:**
*Scheme 1: Synthesis of TCL-1*
*Scheme 2: General Synthesis of TEC Degrader*

**Appendix:**
*$^{1}$H and $^{13}$C NMRs for TEC Degrader Series*

##### *General Materials*

All chemicals, reagents and solvents were purchased from commercial providers and were used as such. All anhydrous solvents were used under nitrogen. The intermediate compounds as well as the target compounds were purified by flash column chromatography using silica gel 60 (0.040-0.063 mm, 230-400 mesh ASTM) and technical grade solvents. The reactions were monitored by thin layer chromatography (TLC) and/or LCMS on Agilent 1260 Infinity II system with Single Quadrupole LC/MS System. $^{1}H$ and $^{13}C$ NMR spectra were obtained on Bruker DRX500-2 spectrometer (500 and 125 MHz for $^{1}H$ and $^{13}C$ NMR respectively). Chemical shifts (δ) are reported in ppm relative to tetramethyl silane (TMS) as internal standard or calibrated using residual $CHCl_3$ peak as δ 7.26 for $^{1}H$ and δ 77.16 for $^{13}C$ NMR. Spin multiplicities for $^{1}H$ NMR are reported as s (singlet), brs (broad singlet), d (doublet), dd (double doublet), t (triplet), q (quartet) and m (multiplet). Coupling constant (*J*) values are reported in hertz (Hz).

##### *Synthesis of TCL-1*

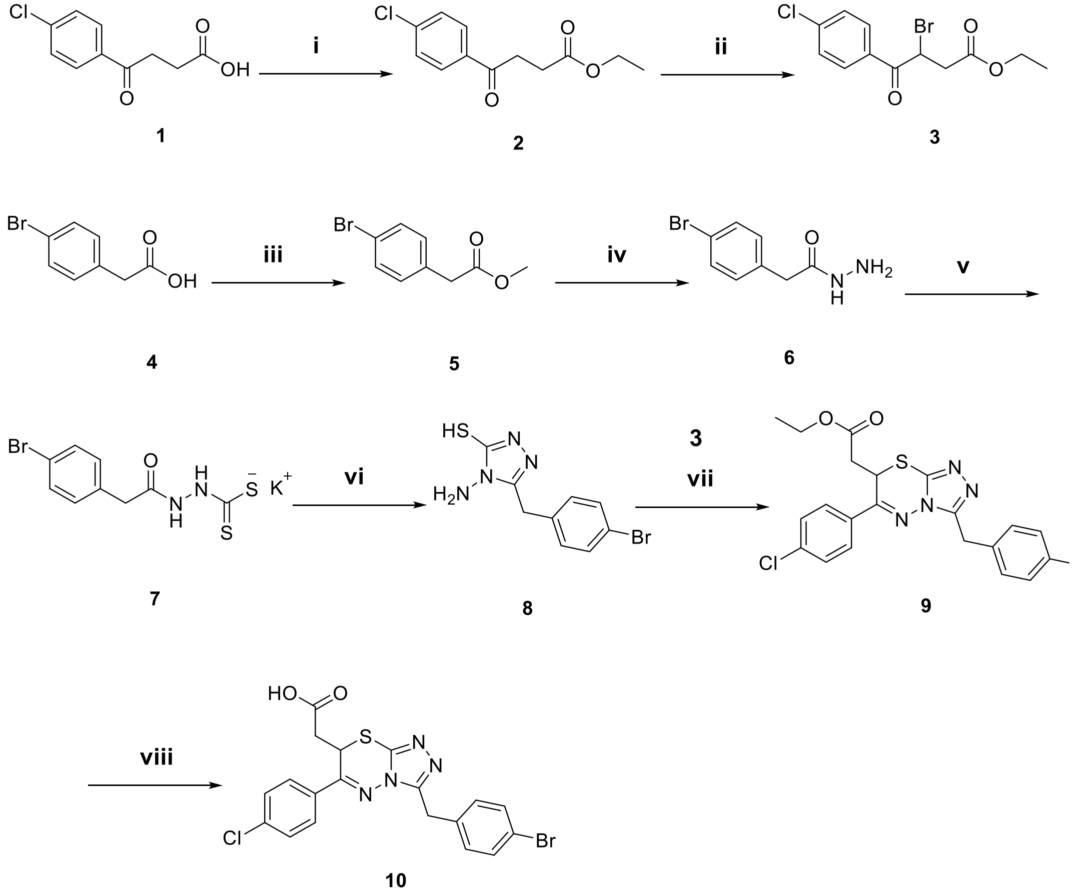


**Scheme 1. Reagents and conditions: i)** EtOH, conc. $H_2SO_4$, 80 °C, 8 h; **ii)** $Br_2$, DCM, 0 °C-rt, 2 h; **iii)** MeOH, conc. $H_2SO_4$, 70 °C, 8 h; **iv)** $NH_2NH_2.H_2O$, MeOH, 70 °C, 12h; **v)** $CS_2$, KOH, MeOH, 0 °C-rt, 4 h; **vi)** $NH_2NH_2.H_2O$, $H_2O$, 100 °C,18h; **vii)** EtOH, 80 °C, 36 h; **viii)** $LiOH.H_2O$, THF:H2O (4:1), rt, 1h.

##### *General synthetic procedures for compounds 2-10*

Step i: To a solution of 4-(4-chlorophenyl)-4-oxobutanoic acid **1** (2.15 g, 10.11 mmol) in ethanol (20 ml), few drops of conc. sulfuric acid were added. The reaction mixture was refluxed for 8 h. The resulting mixture was allowed to cool to rt, diluted with water, neutralized by saturated solution of $NaHCO_3$, extracted with ethyl acetate (3 × 20 mL). The organic layer was dried over anhydrous sodium sulfate and evaporated under vacuum to yield the required ethyl 4-(4-chlorophenyl)-4-oxobutanoate **2** in 95% yield.

Step ii: To a solution of ethyl 4-(4-chlorophenyl)-4-oxobutanoate **2** (773.1 g, 3.21 mmol) in dichloromethane (10 ml) was added bromine (1.1 eq) under ice-cooling, and after stirring at the same temperature for 30 minutes, the reaction mixture was warmed to room temperature and was stirred for one hour. The reaction mixture was poured into ice-water, and DCM. The organic layer was collected, washed with water and brine, dried over anhydrous sodium sulfate and the solvent was removed under vacuum to obtain ethyl 3-bromo-4-(4-chlorophenyl)-4-oxobutanoate **3** in 85% yield.

Step iii: To a solution of 2-(4-bromophenyl)acetic acid 4 (2.01 g, 9.34 mmol) in methanol (20 ml), few drops of conc. sulfuric acid were added. The reaction mixture was refluxed for 8 h. The resulting mixture was allowed to cool to rt, diluted with water, neutralized by saturated solution of $NaHCO_3$, extracted with ethyl acetate (3 × 20 mL). The organic layer was dried over anhydrous sodium sulfate and evaporated under vacuum to yield the desired ethyl 2-(4-bromophenyl)acetate **5** in 97% yield.

Step iv: The respective ester **5** (2.12 g, 9.25 mmol) were dissolved in methanol (50 mL). Hydrazine mono hydrate (4 eq) was added slowly, and the reaction mixture was refluxed for 12 h. After reaction completion, the mixture was concentrated under reduced pressure. The resulting crude solid was filtered, and crystallized from aqueous ethanol to obtain 2-(4-bromophenyl)acetohydrazide **6** in 65% yield.

Step v: The hydrazide compound **6** (1.31 g, 5.75 mol) was treated with a solution of potassium hydroxide (12.5 eq) in methanol (35 mL) at 0°C under stirring. Carbon disulfide (12.5 eq) was added dropwise, and the reaction mixture was stirred for 4 h at 0°C. The potassium dithiocarbazinate **7** product formed was filtered, washed with chilled diethyl ether, and dried at room temperature.

Step vi: The potassium dithiocarbazinate compound **7** was dissolved in distilled water, hydrazine mono hydrate (4 eq) was added, and the reaction mixture was refluxed overnight. Hydrogen sulfide gas evolved, and the reaction mixture turned green. After reaction completion, the mixture was poured in ice and neutralized with concentrated hydrochloric acid. White precipitate was formed, filtered, washed with cold water, and crystallized from methanol to afford 4-amino-5-(4-bromobenzyl)-4H-1,2,4-triazole-3-thiol **8** in 60% yield.

Step vii: A mixture of 4-amino-5-(4-bromobenzyl)-4H-1,2,4-triazole-3-thiol **8** (0.51 g, 1.76 mol) and ethyl 3-bromo-4-(4-chlorophenyl)-4-oxobutanoate **3** (1.1 eq) was dissolved in ethanol (partially dissolved). The reaction mixture was refluxed for 36 h. After the completion of the reaction, the reaction mixture was extracted by ethyl acetate, washed with brine, dried over anhydrous sodium sulfate and evaporated under vacuum. The final product was purified by silica gel (10% MeOH in DCM elution solvent) to yield ethyl 2-(3-(4-bromobenzyl)-6-(4-chlorophenyl)-7H-[1,2,4]triazolo[3,4-*b*][1,3,4]thiadiazin-7-yl)acetate **9** in 60% yield.

**ethyl 2-(3-(4-bromobenzyl)-6-(4-chlorophenyl)-7H-[1,2,4]triazolo[3,4-*b*][1,3,4]thiadiazin-7-yl)acetate** (**9**): **Yield:** 60%. $^1$H NMR (400 MHz, $CDCl_3$) δ 7.77 (d, *J* = 7.77 Hz, 2H, Ar-H), 7.48 (d, *J* = 7.47 Hz, 2H, Ar-H), 7.4 (d, *J* = 7.41 Hz, 2H, Ar-H), 7.21 (m, 2H, Ar-H), 4.76 (dd, *J* = 4.76 Hz, 1H, CH), 4.32 (m, 2H, $CH_2$), 4.12 (m, 2H, $CH_2$), 2.59 (m, 2H, C**H**$_2$CH$_3$), 1.21 (s, 3H, CH$_2$C**H**$_3$). $^{13}$C NMR (100 MHz, $CDCl_3$) δ 168.42 (C=O), 153.95 (Ar-C), 138.81 (Ar-C), 134.21 (Ar-C), 131.47 (Ar-C), 130.45 (Ar-C), 129.57 (Ar-C), 128.39 (Ar-C), 121.07 (Ar-C), 61.87 (**C**H$_2$CH$_3$), 36.95 (CH), 32.76 ($CH_2$), 30.4 ($CH_2$), 13.93 (CH$_2$**C**H$_3$). LC/MS 507.1 $(M+H)^+$.

Step viii: To a solution of ethyl 2-(3-(4-bromobenzyl)-6-(4-chlorophenyl)-7H-[1,2,4]triazolo[3,4-*b*][1,3,4]thiadiazin-7-yl)acetate **9** (0.52 g, 1.03 mmol) in THF (4 ml), a solution of LiOH.$H_2O$ in water (1 ml) was added at 0 °C. The reaction mixture was warmed to room temperature and was stirred for one hour. After the completion of the reaction, the reaction mixture was neutralized by 1M HCl, extracted by DCM, washed with brine, dried over anhydrous sodium sulfate, and evaporated under vacuum. The final product was purified by silica gel (6% MeOH in DCM elution solvent) to yield the targeted 2-(3-(4-bromobenzyl)-6-(4-chlorophenyl)-7H-[1,2,4]triazolo[3,4-b][1,3,4]thiadiazin-7-yl)acetic acid **10** in 50% yield.

**2-(3-(4-bromobenzyl)-6-(4-chlorophenyl)-7H-[1,2,4]triazolo[3,4-b][1,3,4]thiadiazin-7-yl)acetic acid (10): Yield:** 55%. $^1$H NMR (500 MHz, DMSO *d6*) δ 7.96 (m, 2H), 7.60 (dd, *J* = 8.6, 1.5 Hz, 2H), 7.47 (m, 2H), 7.25 (m, 2H), 5.13 (ddd, *J* = 10.1, 4.8, 1.3 Hz, 1H), 4.27 (m, 2H), 2.66 (ddd, *J* = 16.6, 4.9, 1.4 Hz, 1H), 2.46 (m, 2H). $^{13}$C NMR (126 MHz, DMSO *d6*) δ 170.25, 155.14, 152.49, 138.99, 137.25, 135.79, 131.73, 131.60, 131.42, 129.80,

129.52, 120.27, 40.10, 39.93, 39.77, 39.60, 39.43, 39.27, 36.76, 33.30, 29.82. LCMS (ESI) m/z $C_{19}H_{14}BrClN_4O_2S$: calcd, 477.7; found, 479.0 $[M + H]^+$.

***Synthesis of TCL1 Degraders:***

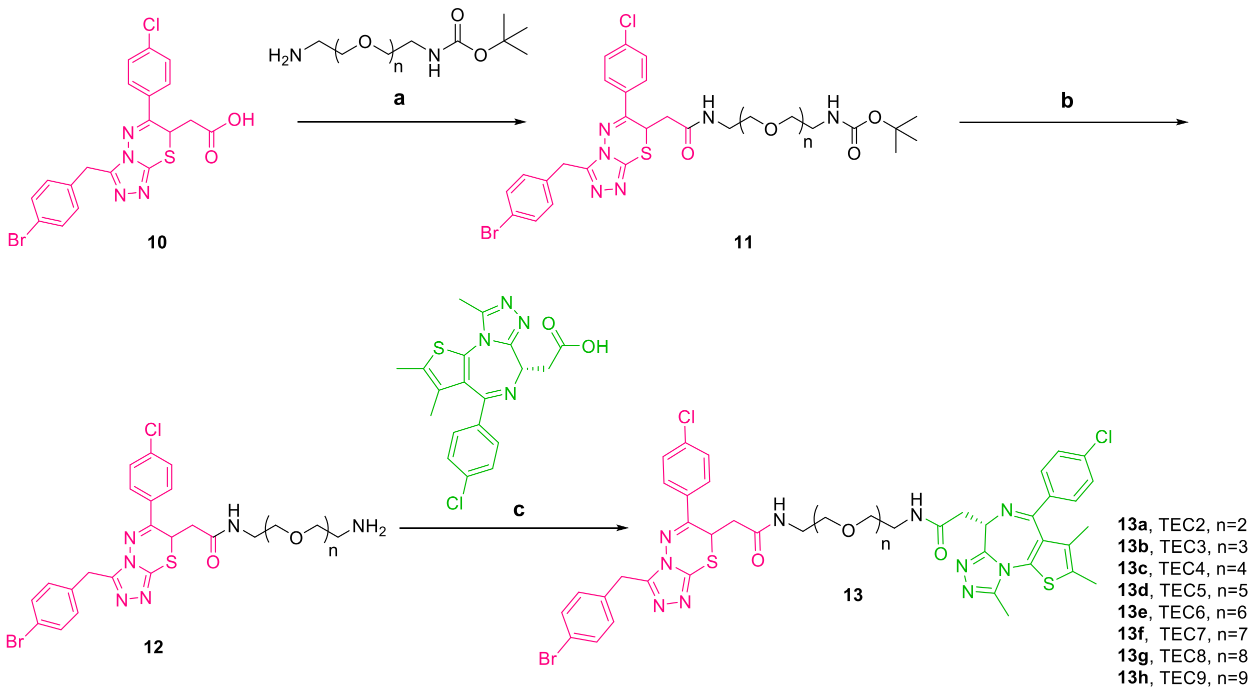


**Scheme 2. Reagents and Conditions: a)** HOBt.$H_2O$, EDC, DIPEA, THF, rt,12h; **b)** TFA:DCM (1:1), rt, 12h; **c)** HOBt.$H_2O$, EDC, DIPEA, THF, rt,12h.

***General synthetic procedures for TEC2-9:***

Step a: To a solution of 2-(3-(4-bromobenzyl)-6-(4-chlorophenyl)-7H-[1,2,4]triazolo[3,4-b][1,3,4]thiadiazin-7-yl)acetic acid, TCL-1 compound **10** (1 eq.), HOBt.$H_2O$ (1.5 eq.), and EDC (1.5 eq.) in THF, DIPEA (6 eq.) was added, and the reaction mixture was stirred for 15 minutes before adding the corresponding BOC protected PEG linkers. After linkers addition, the reaction mixture was stirred at room temperature for 12 h. After reaction completion (TLC monitoring), the reaction mixture was poured into saturated $NaHCO_3$ solution, and extracted with ethyl acetate. The organic layer was washed with water and brine, dried over anhydrous sodium sulfate and the solvent was removed under vacuum. The crude product was purified by silica gel (10% MeOH in DCM elution solvent) to obtain compound **11**. in 50-60% yield.

Step b: To a solution of compound **11** in DCM, equal volume of TFA was added. The reaction mixture was stirred at room temperature for 8 h. After reaction completion (TLC monitoring), the reaction mixture was poured into saturated $NaHCO_3$ solution, and extracted with DCM for 3 times. The organic layers were washed with water and brine, dried over anhydrous sodium sulfate and the solvent was removed under vacuum. Crude product **12** was subjected to the next step without further purification.

Step c: To a solution of (S)-2-(4-(4-chlorophenyl)-2,3,9-trimethyl-6H-thieno[3,2-f][1,2,4]triazolo[4,3-a][1,4]diazepin-6-yl)acetic acid **JQ1** (1 eq.), HOBt.$H_2O$ (1.5 eq.), and EDC (1.5 eq.) in THF, DIPEA (6 eq.) was added, and the reaction mixture was stirred for 15 minutes before adding the compound **12**. By addition of **3**, the reaction mixture was stirred at room temperature for 12 h. After reaction completion (TLC monitoring), the reaction mixture was poured into saturated $NaHCO_3$ solution, and extracted with ethyl acetate. The organic layer was washed with water and brine, dried over anhydrous sodium sulfate and the solvent was removed under vacuum. The crude product was purified by silica gel (8% MeOH in DCM elution solvent) to obtain the target compound **13** (**TEC2-9**) in 40-50% yield.

**2-(3-(4-bromobenzyl)-6-(4-chlorophenyl)-7H-[1,2,4]triazolo[3,4-b][1,3,4]thiadiazin-7-yl)-N-(2-(2-(2-(2-((S)-4-(4-chlorophenyl)-2,3,9-trimethyl-6H-thieno[3,2-f][1,2,4]triazolo[4,3-a][1,4]diazepin-6-yl)acetamido)ethoxy)ethoxy)ethyl)acetamide (TEC2):** $^{1}$H NMR (500 MHz, $CDCl_3$) δ 7.86 – 7.77 (m, 2H), 7.40 (ddd, *J* = 9.1, 5.1, 2.2 Hz, 4H), 7.27 (d, *J* = 7.6 Hz, 3H), 7.15 – 7.04 (m, 2H), 4.93 (dd, *J* = 10.1, 4.1 Hz, 1H), 4.67 (s, 1H), 4.30 (s, 1H), 4.1-3.7 (m, 1H), 3.60 (s, 4H), 3.56 (ddd, *J* = 9.2, 5.8, 2.7 Hz, 6H), 3.50 – 3.42 (m, 2H), 3.41 – 3.36 (m, 2H), 2.69 (dt, *J* = 18.9, 7.9 Hz, 1H), 2.78-2.65 (m, 1H), 2.57 – 2.46 (m, 3H), 2.38 (s, 3H), 2.05 (s, 1H), 1.64 (s, 3H), 1.35-1.2 (m, 3H). $^{13}$C NMR (126 MHz, $CDCl_3$) δ 170.32, 167.63, 163.97, 155.32, 154.79, 152.32, 149.83, 139.44, 138.36, 136.70, 136.31, 134.33, 131.63, 131.51, 131.01, 130.66, 130.46, 130.02, 129.81, 129.33, 128.58, 120.74, 70.30, 70.10, 69.80, 69.68, 69.56, 54.18, 54.09, 39.76, 39.67, 39.47, 39.30, 38.85, 38.61, 38.31, 38.24, 33.62, 33.49, 30.32, 14.32, 13.03, 11.73. LCMS (ESI) *m/z* $C_{44}H_{43}BrCl_2N_{10}O_4S_2$: calcd, 990.8; found, 991.2 $[M + H]^+$.

**2-(3-(4-bromobenzyl)-6-(4-chlorophenyl)-7H-[1,2,4]triazolo[3,4-b][1,3,4]thiadiazin-7-yl)-N-(1-((S)-4-(4-chlorophenyl)-2,3,9-trimethyl-6H-thieno[3,2-f][1,2,4]triazolo[4,3-a][1,4]diazepin-6-yl)-2-oxo-6,9,12-trioxa-3-azatetradecan-14-yl)acetamide (TEC3):** $^{1}$H NMR (500 MHz, $CDCl_3$) δ 8.04 (s, 1H), 7.84 (dd, *J* = 8.6, 4.0 Hz, 2H), 7.58 (d, *J* = 20.0 Hz, 1H), 7.45 – 7.35 (m, 4H), 7.33 (dd, *J* = 8.3, 2.8 Hz, 2H), 7.30 – 7.24 (m, 2H), 7.16 (dd, *J* = 8.1, 5.1 Hz, 2H), 5.01 – 4.87 (m, 1H), 4.72 (m, 1H), 4.27 (q, *J* = 10.8 Hz, 2H), 3.64 (d, *J* = 6.5 Hz, 3H), 3.61 (d, *J* = 4.1 Hz, 6H), 3.58 – 3.50 (m, 4H), 3.43 (dd, *J* = 12.5, 6.8 Hz, 5H), 2.89 – 2.67 (m, 1H), 2.61 (d, *J* = 10.0 Hz, 3H), 2.54 (dd, *J* = 15.3, 4.0 Hz, 1H), 2.40 (d, *J* = 2.4 Hz, 3H), 1.67 (s, 3H). $^{13}$C NMR (126 MHz, $CDCl_3$) δ 169.15, 167.62, 139.62, 139.46, 133.66, 133.54, 132.07, 132.01, 131.73, 131.29, 131.07, 130.14, 130.04, 129.67, 129.19, 129.07, 128.90, 121.64, 70.51, 70.17, 70.08, 69.54, 69.37, 53.07, 39.65, 39.50, 39.42, 38.77, 36.62, 33.86, 33.62, 30.07, 14.53, 13.35, 11.97. LCMS (ESI) m/z $C_{46}H_{47}BrCl_2N_{10}O_5S_2$: calcd, 1034.87; found, 1035.2 $[M + H]^+$.

**2-(3-(4-bromobenzyl)-6-(4-chlorophenyl)-7H-[1,2,4]triazolo[3,4-b][1,3,4]thiadiazin-7-yl)-N-(1-((S)-4-(4-chlorophenyl)-2,3,9-trimethyl-6H-thieno[3,2-f][1,2,4]triazolo[4,3-a][1,4]diazepin-6-yl)-2-oxo-6,9,12,15-tetraoxa-3-azaheptadecan-17-yl)acetamide (TEC4):** $^{1}$H NMR (400 MHz, $CDCl_3$) δ 7.83 (dd, *J* = 8.8, 2.7 Hz, 2H), 7.43 (dt, *J* = 8.8, 2.5 Hz, 3H), 7.41 – 7.35 (m, 3H), 7.34 – 7.28 (m, 2H), 7.18 (dd, *J* = 8.2, 4.8 Hz, 2H), 4.93 (d, *J* = 9.2 Hz, 1H), 4.77 (dt, *J* = 25.9, 6.8 Hz, 1H), 4.26 (s, 2H), 3.75 – 3.51 (m, 18H), 3.45 (d, *J* = 9.8 Hz, 6H), 2.69 (d, *J* = 14.4 Hz, 4H), 2.41 (d, *J* = 3.0 Hz, 3H), 1.67 (d, *J* = 7.2 Hz, 3H).$^{13}$C NMR (101 MHz, $CDCl_3$) δ 170.48, 167.77, 138.77, 132.06, 131.98, 130.94, 130.84, 130.36, 129.69, 128.96, 128.93, 128.83, 121.24, 77.60, 77.29, 76.97, 70.73, 70.65, 70.56, 70.41, 70.03, 69.70, 54.06, 39.79, 39.65, 38.64, 33.86, 30.65, 14.63, 13.38, 11.86. LCMS (ESI) *m/z* $C_{48}H_{51}BrCl_2N_{10}O_6S_2$: calcd, 1078.92; found, 1079.2 $[M + H]^+$.

**2-(3-(4-bromobenzyl)-6-(4-chlorophenyl)-7H-[1,2,4]triazolo[3,4-b][1,3,4]thiadiazin-7-yl)-N-(1-((S)-4-(4-chlorophenyl)-2,3,9-trimethyl-6H-thieno[3,2-f][1,2,4]triazolo[4,3-a][1,4]diazepin-6-yl)-2-oxo-6,9,12,15,18-pentaoxa-3-azaicosan-20-yl)acetamide (TEC5):** $^{1}$H NMR (500 MHz, $CDCl_3$) δ 7.85 (d, *J* = 8.7 Hz, 2H), 7.47 – 7.42 (m, 2H), 7.38 (td, *J* = 8.4, 6.0 Hz, 4H), 7.32 – 7.28 (m, 2H), 7.18 (dd, *J* = 8.5, 2.5 Hz, 2H), 4.96 (dd, *J* = 10.1, 3.9 Hz, 1H), 4.68 (q, *J* = 7.5 Hz, 1H), 4.25 (d, *J* = 2.9 Hz, 2H), 3.57 (d, *J* = 5.2 Hz, 11H), 3.54 (dt, *J* = 7.9, 2.4 Hz, 9H), 3.49 – 3.27 (m, 7H), 2.65 (d, *J* = 7.4 Hz, 3H), 2.56 – 2.45 (td, J = 14.6, 3.9 Hz, 1H), 2.39 (s, 3H), 1.66 (s, 3H). $^{13}$C NMR (126 MHz, $CDCl_3$) δ 170.35, 167.28, 163.82, 162.08, 155.49, 154.76, 152.31, 149.86, 138.47, 131.65, 130.89, 130.52, 129.85, 129.41, 128.55, 120.89, 70.40, 70.35, 70.31, 70.22, 69.71, 69.41, 54.05, 39.75, 39.32, 38.54, 38.36, 33.31, 30.33, 14.34, 13.05, 11.67. LCMS (ESI) *m/z* $C_{50}H_{55}BrCl_2N_{10}O_7S_2$: calcd, 1122.98; found, 1123.2 $[M + H]^+$.

**2-(3-(4-bromobenzyl)-6-(4-chlorophenyl)-7H-[1,2,4]triazolo[3,4-b][1,3,4]thiadiazin-7-yl)-N-(1-((R)-4-(4-chlorophenyl)-2,3,9-trimethyl-6H-thieno[3,2-f][1,2,4]triazolo[4,3-a][1,4]diazepin-6-yl)-2-oxo-6,9,12,15,18,21-hexaoxa-3-azatricosan-23-yl)acetamide (TEC6):** $^{1}$H NMR (500 MHz, $CDCl_3$) δ 7.86 (m, 2H), 7.45 (dd, *J* = 8.7, 2.2 Hz, 2H), 7.43 – 7.39 (m, 2H), 7.37 (d, *J* = 8.3 Hz, 2H), 7.32 (d, *J* = 8.5 Hz, 2H), 7.21 (dd, *J* = 8.3, 4.4 Hz, 2H), 4.95 (dt, *J* = 9.8, 4.5 Hz, 1H), 4.74 (q, *J* = 6.5 Hz, 1H), 4.37 – 4.18 (m, 2H), 3.72 – 3.62 (m, 7H), 3.62 – 3.58 (m, 8H), 3.57 (d, *J* = 3.2 Hz, 3H), 3.54 (ddt, *J* = 12.5, 8.2, 3.2 Hz, 6H), 3.48 (d, *J* = 5.8 Hz, 10H), 2.73 (s, 3H), 2.41 (s, 3H), 1.67 (d, *J* = 3.2 Hz, 3H). $^{13}$C NMR (126 MHz, $CDCl_3$) δ 131.75, 131.70, 130.69, 130.58, 130.01, 129.43, 128.64, 70.37, 69.66, 39.73, 39.35, 14.35, 13.09. LCMS (ESI) *m/z* $C_{52}H_{59}BrCl_2N_{10}O_8S_2$: calcd, 1167.03; found, 1167.2 $[M + H]^+$.

**2-(3-(4-bromobenzyl)-6-(4-chlorophenyl)-7H-[1,2,4]triazolo[3,4-b][1,3,4]thiadiazin-7-yl)-N-(1-((S)-4-(4-chlorophenyl)-2,3,9-trimethyl-6H-thieno[3,2-f][1,2,4]triazolo[4,3-a][1,4]diazepin-6-yl)-2-oxo-6,9,12,15,18,21,24-heptaoxa-3-azahexacosan-26-yl)acetamide (TEC7):** $^{1}$H NMR (500 MHz, $CDCl_3$) δ 7.92 – 7.77 (m, 2H), 7.45 – 7.41 (m, 2H), 7.41 – 7.35 (m, 4H), 7.29 (d, *J* = 8.3 Hz, 2H), 7.17 (dd, *J* = 8.3, 3.2 Hz, 2H), 4.93 (ddd, *J* = 10.2, 4.0, 1.8 Hz, 1H), 4.65 (t, *J* = 6.9 Hz, 1H), 4.25 (s, 2H), 3.61 (s, 10H), 3.58 – 3.54 (m, 10H), 3.50 (ddd, *J* = 8.1, 6.0, 3.7 Hz, 9H), 3.47 – 3.41 (m, 5H), 3.40 – 3.31 (m, 2H), 2.64 (d, *J* = 1.8 Hz, 3H), 2.38 (s, 3H), 1.64 (s, 3H). $^{13}$C NMR (126 MHz, $CDCl_3$) δ 170.35, 167.23, 163.79, 159.36, 155.48, 154.65, 149.82, 138.47, 136.67, 136.38, 134.43, 131.82, 131.69, 130.85, 130.55, 129.82, 129.42, 128.55, 120.91,70.41, 70.33, 70.23, 70.18, 69.69, 69.41, 54.10, 39.73, 39.29, 38.66, 38.24, 33.31, 30.31, 14.33, 13.03, 11.69. LCMS (ESI) *m/z* $C_{54}H_{63}BrCl_2N_{10}O_9S_2$: calcd, 1211.08; found, 1211.2 $[M + H]^+$.

**2-(3-(4-bromobenzyl)-6-(4-chlorophenyl)-7H-[1,2,4]triazolo[3,4-b][1,3,4]thiadiazin-7-yl)-N-(1-((R)-4-(4-chlorophenyl)-2,3,9-trimethyl-6H-thieno[3,2-f][1,2,4]triazolo[4,3-a][1,4]diazepin-6-yl)-2-oxo-6,9,12,15,18,21,24,27-octaoxa-3-azanonacosan-29-yl)acetamide (TEC8):** $^{1}$H NMR (500 MHz, $CDCl_3$) δ 7.92 – 7.78 (m, 2H), 7.51 – 7.43 (m, 2H), 7.43 – 7.35 (m, 4H), 7.30 (d, *J* = 8.0 Hz, 2H), 7.21 – 7.20 (m, 2H), 7.19 (d, *J* = 7.19 Hz, 1H),7.06 (d, *J* = 6.2 Hz, 1H), 5.01 – 4.87 (m, 1H), 4.66 (t, *J* = 6.9 Hz, 1H), 4.27 (s, 2H), 3.66 – 3.62 (m, 8H), 3.61 – 3.57 (m, 10H), 3.56 – 3.51 (m, 10H), 3.51 – 3.41 (m, 9H), 3.41 – 3.35 (m, 3H), 2.66 (s, 3H), 2.39 (s, 3H), 1.65 (s, 3H), 1.24 (s, 1H). $^{13}$C NMR (126 MHz, $CDCl_3$) δ 170.35, 167.17, 163.80, 155.49, 154.59, 152.37, 149.81, 139.26, 138.52, 136.69, 136.40, 134.46, 131.71, 130.88, 130.52, 129.82, 129.45, 128.58, 120.94, 70.44, 70.39, 70.32, 70.25, 70.18, 69.69, 69.43, 54.14, 39.77, 39.31, 38.76, 38.23, 33.30, 30.35, 14.33, 13.03, 11.72. LCMS (ESI) *m/z* $C_{56}H_{67}BrC_{l2}N_{10}O_{10}S_2$: calcd, 1255.1; found, 1255.2 $[M + H]^+$.

**2-(3-(4-bromobenzyl)-6-(4-chlorophenyl)-7H-[1,2,4]triazolo[3,4-b][1,3,4]thiadiazin-7-yl)-N-(1-((S)-4-(4-chlorophenyl)-2,3,9-trimethyl-6H-thieno[3,2-f][1,2,4]triazolo[4,3-a][1,4]diazepin-6-yl)-2-oxo-6,9,12,15,18,21,24,27,30-nonaoxa-3-azadotriacontan-32-yl)acetamide (TEC9):** $^{1}$H NMR (500 MHz, $CDCl_3$) δ 7.86 (dd, *J* = 8.6, 2.1 Hz, 2H), 7.46 (d, *J* = 8.5 Hz, 2H), 7.40 (ddd, *J* = 9.8, 5.3, 2.2 Hz, 4H), 7.31 (dd, *J* = 8.3, 3.2 Hz, 2H), 7.21 (d, *J* = 8.0 Hz, 2H), 4.94 (d, *J* = 9.5 Hz, 1H), 4.77 – 4.60 (m, 1H), 4.29 (s, 2H), 3.65 (s, 10H), 3.62 (s, 5H), 3.58 (d, *J* = 5.7 Hz, 10H), 3.54 (d, *J* = 7.9 Hz, 10H), 3.49 (dd, *J* = 15.1, 6.0 Hz, 9H), 3.40 (dd, *J* = 15.5, 6.8 Hz, 3H), 2.71 (s, 3H), 1.66 (s, 3H). $^{13}$C NMR (126 MHz, $CDCl_3$) δ 169.81, 167.35, 164.92, 156.27, 155.07, 150.53, 139.04, 137.71, 135.25, 133.69, 133.49, 132.81, 131.93, 131.53, 130.94, 130.46, 129.62, 128.82, 121.31, 70.58, 70.55, 70.52, 70.47, 70.41, 70.33, 53.71, 41.01, 39.74, 39.49, 38.48, 37.77, 33.50, 30.32, 14.48, 13.29, 11.66. LCMS (ESI) *m/z* $C_{58}H_{71}BrCl_2N_{10}O_{11}S_2$: calcd, 1299.19; found, 1299.2 $[M + H]^+$.

***General Cell Culture:***

HEK-293T (ATCC®) (CRL-3216TM) were grown in DMEM (Dulbecco's Modified Eagle's Medium, ATCC® 30-2002TM) supplemented with 10% fetal bovine serum (FBS), U87 (HTB-14) were grown in EMEM (Eagle's Minimum Essential Medium, ATCC® 30-2003TM) supplemented with 10% FBS and 1x penicillin/streptomycin; and both Ramos (ATCC® CRL-1596TM) and Raji (ATCC® CCL-86) were grown in RPMI-1640 (ATCC® 30-2001TM) supplemented with 10% FBS. All cells were grown in a humidified 37ºC incubator at 5% $CO_2$. For suspension cells (Ramos and Raji), the cell media was pelleted @ 800 xg for 5 min. For adherent cells (HEK-293T and U87) the media was aspirated, and the flask rinsed with sterile PBS before treating cells with 0.25% (w/v) Trypsin-0.53 mM EDTA solution. The flasks were returned to the incubator for 5 min. Trypsin was quenched with supplemented media, suspension was collected and cells pelleted @ 1000 xg for 5 min. After aspirating media, the cell pellet was resuspended in fresh media with 1% FBS. Cell concentration was determined by Trypan Blue, a hemocytometer and plated accordingly.

***BRD4 Degradation Dosing with Suspension cells (Ramos & Raji)***

Suspension cells were plated in 12 well clear plates at a density of 400,000 cells/well (500 µL per well of a 800,000 cell/mL stock). Cells were then treated with 500 µL of 2x target concentration of compound stocked prepared at an initial 1% DMSO solution in cell media (1% FBS), in triplicate. This dilution makes the final 1 mL solution 0.5% DMSO overall and 1x target concentration of compound. Cell plate was placed in incubator for 18 hrs. After 18 hours, cells were collected in 1.5 mL Eppendorf tubes and spun down at 800 xg for 5 mins. Media aspirated and pellet resuspended in RIPA lysis buffer supplemented with 1x HALT protease inhibitor and allowed to lyse for 10 mins before centrifuging at 14,000 xg for 20 mins.

##### *BRD4 Degradation Dosing with Adherent cells (HEK & U87)*

Adherent cells were plated in 12 well clear plates at a density of 400,000 cells/well (1000 µL per well of a 400,000 cell/mL stock). Cells were allowed to adhere in incubator overnight. Media removed cells were treated with 1000 µL of 1x target concentration of compound stocked prepared at 0.5% DMSO solution in cell media (1% FBS), in triplicate. Cell plate was placed in incubator for 18 hrs. After 18 hours, media was removed, and RIPA lysis buffer supplemented with HALT protease inhibitor was added to each well. Plates were lightly shaken for 10 mins while cells lysed. Lysed cells were collected in 1.5 mL Eppendorf tubes and centrifuged at 14000 xg for 20 mins.

##### *BRD4 Degradation Western Blot*

After spinning lysates at 14000 xg for 20 mins, supernatant was collected, and pellet discarded. Protein concentration was quantitated by Pierce™ BCA Assay Kit (Thermo Fisher Cat. No 23227). Lysates were normalized, approximately 30-40 µg of protein. Lysates were ran under denaturing conditions on 10-well Mini-PROTEAN TGX Precase Gels (Biorad cat. No. 4561094). After SDS-PAGE, gels were cut so that MW bands 100 kD and higher are separated from 75 kD and below. The high molecular weight bands containing BRD4 were transferred to nitrocellulose (0.2 µm) with Biorad Trans-Blot Turbo on the high MW transfer setting for 10 min. The lower MW bands containing tubulin and rpn-13 were transferred to nitrocellulose (0.2 µm) with Biorad Trans-Blot Turbo on turbo setting for 7 min. After transferring, membranes were washed with PBS (5 mins x3). High MW membranes were blocked with 5% milk in PBS for 1 hour at RT. Lower MW bands were blocked with Intercept® (PBS) Blocking Buffer (LI-COR Cat. No. 927-70001) for 1 hr at RT. Primaries, Tubulin (Thermofisher, T5168, mouse, 1:2000) ADRM1 (ABCAM, ab157185, rabbit 1:1000), Brd4 (Thermofisher, A301-985-A-T, Rabbit, 1:1000) were added and incubated with membranes at 4 deg C overnight. The next day primaries were removed, and membranes washed with PBS (10 mins x3). Secondaries (tubulin= IRDYE® 800CW Goat anti-mouse 800 1:10000, ADRM1= IRDye® 680 Goat Anti-Rabbit 1:10000, BRD4= Invitrogen Goat anti-mouse IgG (H+L) HRP 20 ng/mL) were added to membranes and incubated at RT for 1 hr. Secondaries removed, and blots were washed with PBST (0.01% Tween20) 3x 10 mins. Lower MW bands were imaged on Azure 600 Western Blot Imaging System on the fluorescent label setting for 800CW and 680RD. Higher MW membranes were incubated with ThermoFisher Scientific SuperSignal West Pico PLUS chemiluminescent substrate for 5 mins before imaging on AZURE 600 on the chemiluminescent setting. Bands normalized to tubulin to determine amount of BRD4 degraded.

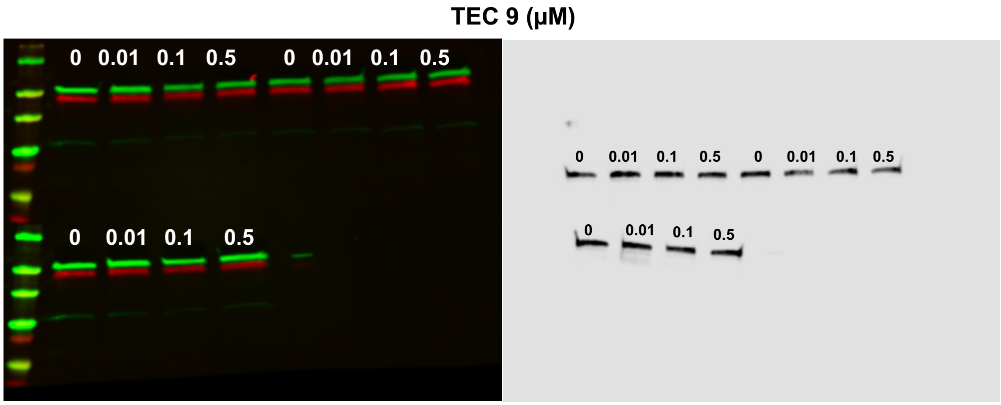


**Figure S1.** Representative western blot for degradation studies. All degraders and cell lines were handled in same fashion. All signals were assessed on Licor Image Studio software, and bands normalized to Tubulin (Green). Rpn-13 was used to demonstrate degraders are not degrading Rpn-13 as well (Red). Brd4 bands were cute and transferred on different setting and blotting with HRP antibody, and normalized to respective DMSO values, (Black bands on right).

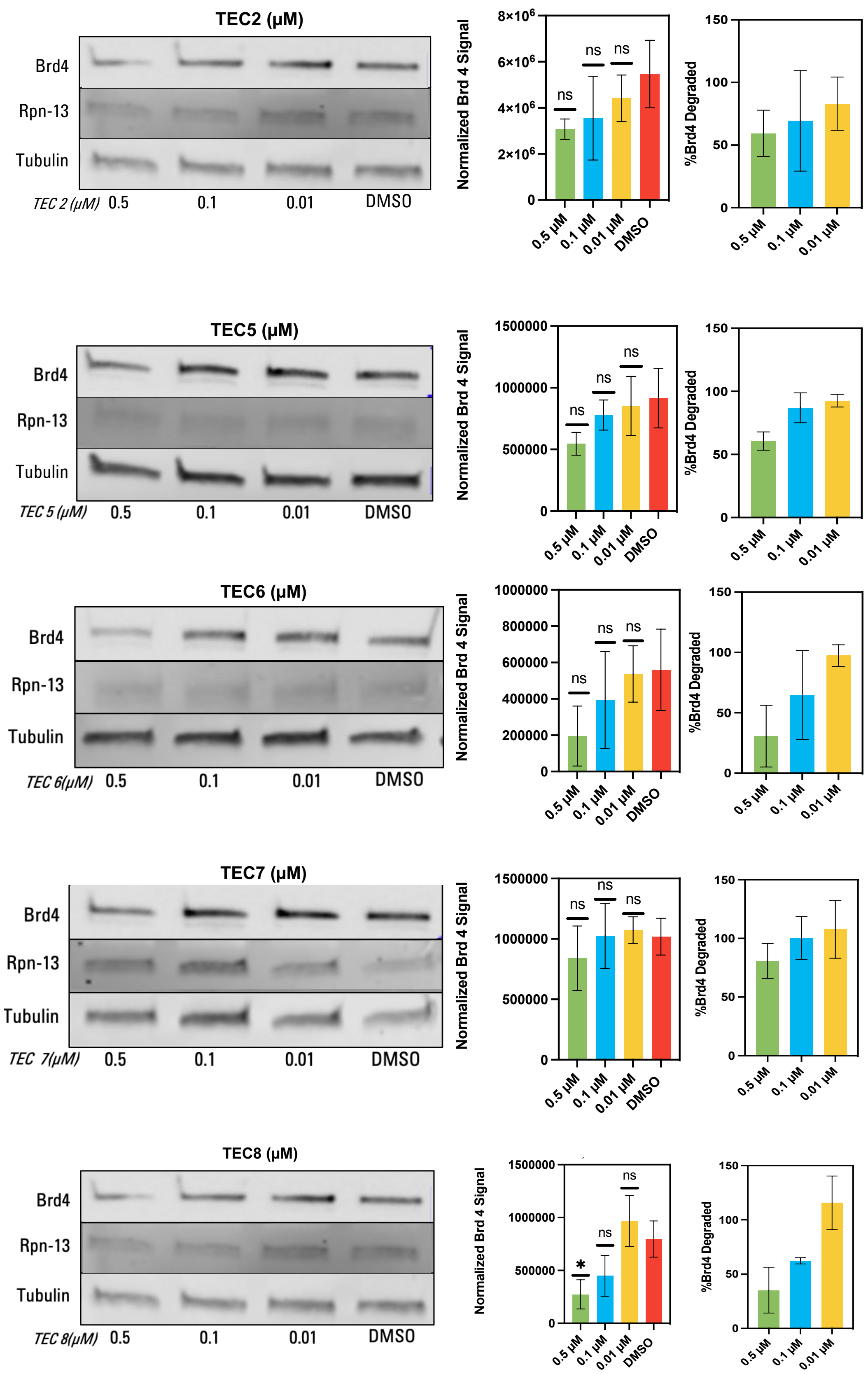


**Figure S2.** Western blots for other degraders (TEC2,TEC5,TEC6,TEC7 and TEC8) in Ramos cells. Degradation was done for 18 hrs before subjecting cells to western blot. Signals were normalized to tubulin, and percent BRD4 remaining calculated with DMSO as 100 %.

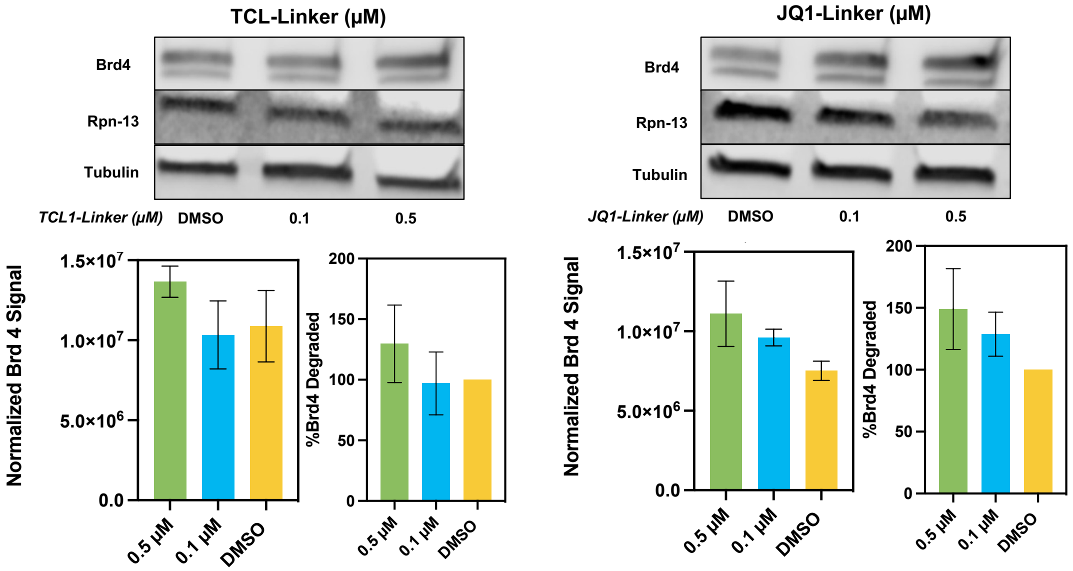


**Figure S3.** BRD4 degradation with Linker controls TCL-Linker (Left) and JQ1-Linker (Right)

##### *Proteasome dependent degradation*

Ramos cells were plated in 12 well plate as mentioned before. Mg-132 was prepared at 25 nM and TEC4 was prepared at 0.01 µM. Cells were dosed with either MG-132, MG-132 and TEC4, or DMSO for 18 hrs. Cells were then subjected to the same western blotting analysis a s stated before.

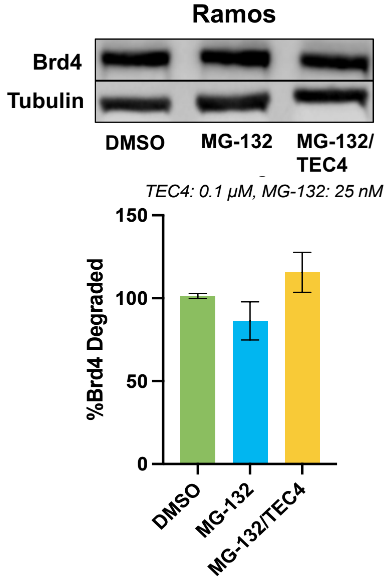


**Figure S4.** MG132/TEC4 dosing to ensure degradation is proteasome dependent.

##### *ADRM1 siRNA*

U87 cells were plated in 24 well plated at 200,000 cells/well (500 µL of a 400,000 cell/mL stock) and placed in incubator to adhere overnight. The following day cells were treated following Dharmafect™ Transfection Reagents -siRNA Transfection Protocol. ON-TARGETplus siRNA (Cat ID:L-012340-01-0020) ADRM1 SMARTPool 20 nmol was purchased from horizon and resuspended in 1x buffer (5x buffer: 300 mM KCl, 30 mM HEPES pH: 7.5, 1.0 mM $MgCl_2$) to get to 20 µM stock. siRNA was prepared at a range of concentrations from 0-100 nM by diluting in serum free media. 0.7 µL of transfection reagent (DharmaFECT 1 Transfection Reagent, Cat ID: T-2001-01) was diluted in 174.3 µL of serum free media and allowed to incubate for 5 mins. siRNA in media was added to the transfection reagent and allowed to incubate for 20 mins at RT. After 20 min 1400 µL of media was added to get to a total volume of 1750 µL total. Contents were added to cells in the 24 well plate and placed in incubator for 48 hrs. Media removed and RIPA buffer added to wells to lyse cells. Lysed cells are subjected to western blot procedure as described previously to determine levels of ADRM1.

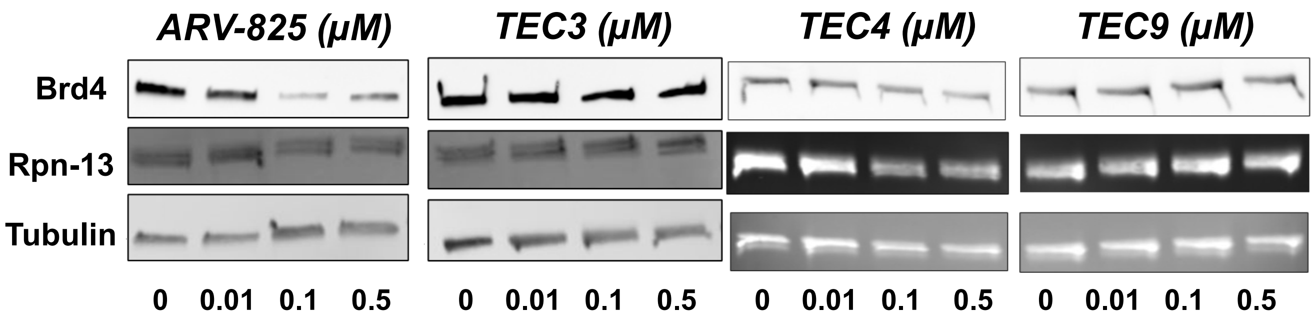


**Figure S5.** HEK293T cell BRD4 degradation with TEC4 and TEC8.

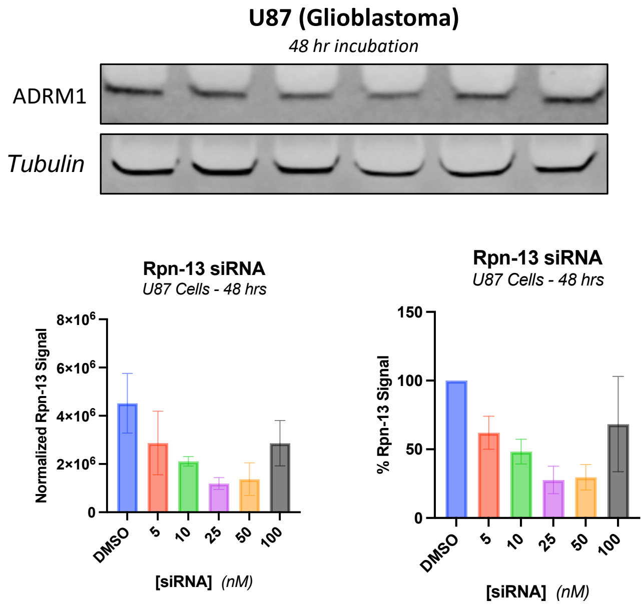


**Figure S6.** siRNA of U87 Cells

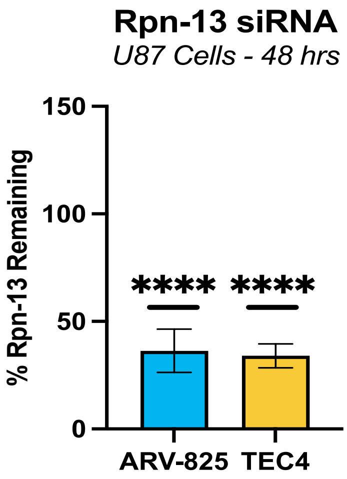


**Figure S7.** Rpn-13 levels remaining after siRNA for 48 hrs and dosing with degraders for 18hrs compared to DMSO control.

##### *ADRM1 siRNA with TEC4 and ARV*

U87 cells were plated in 24 well plated at 200,000 cells/well (500 μL of a 400,000 cell/mL stock) and placed in incubator to adhere overnight. The following day cells were treated following Dharmafect™ Transfection Reagents -siRNA Transfection Protocol. ON-TARGETplus siRNA (Cat ID:L-012340-01-0020) ADRM1 SMARTPool 20 nmol was purchased from horizon and resuspended in 1x buffer (5x buffer: 300 mM KCl, 30 mM HEPES pH: 7.5, 1.0 mM $MgCl_2$) to get to 20 μM stock. siRNA was prepared at optimal concentration to lower ADRM1 signal (25 nM). 0.7 μL of transfection reage nt (DharmaFECT 1 Transfection Reagent, Cat ID: T-2001-01) was diluted in 174.3 μL of serum free media and allowed to incubate for 5 mins. siRNA in media was added to the transfection reagent and allowed to incubate for 20 mins at RT. After 20 min 1400 μL of media was added to get to a total volume of 1750 μL total. Contents were added to cells in the 24 well plate and placed in incubator for 48 hrs. After 48 hrs, compounds were prepared for degradation as previously mentioned for a total of 500 uL/well. Media removed and replaced with media plus degrader and allowed to incubate overnight. The following day media removed, and cells subjected to western blot analysis as previously described to determine degradation.

##### *TAK-243 E1 Ubiquitin Inhibition*

Ramos cells were plated in 12 well plate as mentioned before. TAK-243 was prepared at 1 nM, TEC4 and ARV-825 was prepared at 100 nM. Cells were dosed with either TAC-243 and TEC 4, ARV-825 and TEC 4, or DMSO for 18 hrs. Cells were then subjected to the same western blotting analysis as stated before.

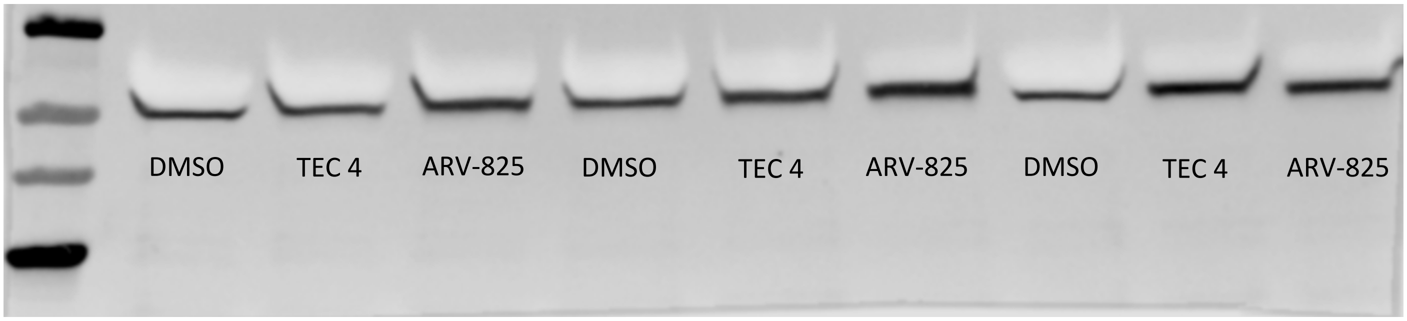


**Figure S8.** Tubulin western blot of TAK-243 dosed cells with ARV-825 and TEC 4

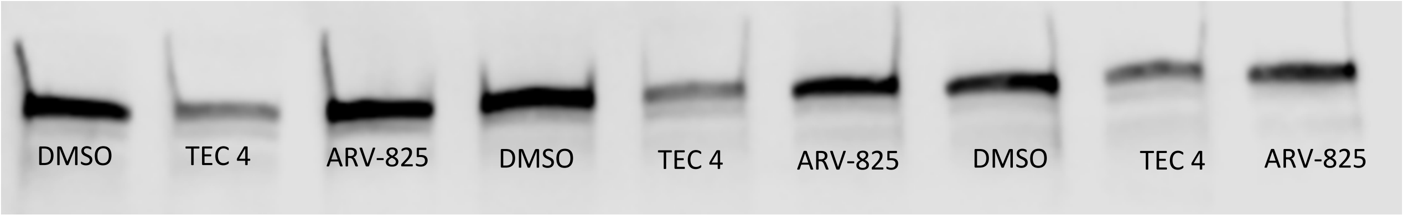


**Figure S9.** BRD4 western blot of TAK-243 dosed cells with ARV-825 and TEC 4

##### *Cellular Toxicity with Suspension Cells (Ramos & Raji)*

Suspension cells were plated in 96 well opaque plates at a density of 10,000 cells/well (50 µL per well of a 200,000 cells/mL stock). 200 µL of PBS was added to surrounding wells. Cells were then treated with 50 µL of 2X target concentration of compound stocks prepared at an initial 1% DMSO solution in cell media (1% FBS), in

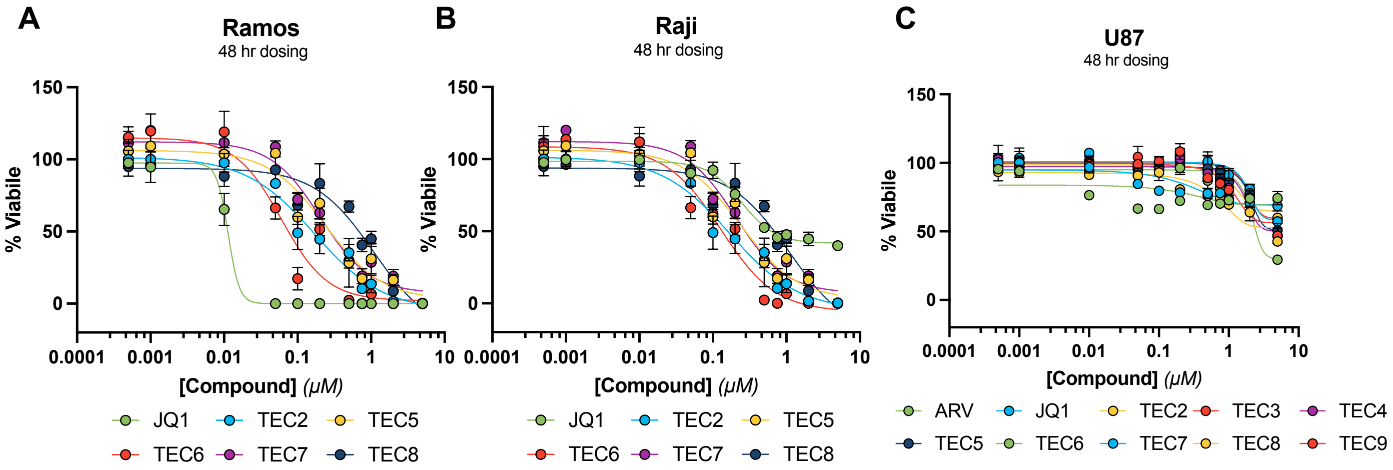


**Figure S10. A)** Ramos cell viability, **B)** Raji Cell viability, **C)** U87 Cell viability. All toxicity readings done at 48 hrs.

triplicate. This dilution makes the final 100 µL solution 0.5% DMSO overall at 1x the target concentration of compound. Cell plate was placed into the incubator for 48 hours. After 48 hours, cell plate was removed and brought to r.t. for 30 mins. 90 µL of Cell Titer-Glo® luminescent reagent was added to each well. Plate was gently shaken for 2 minutes. Plate was then protected from light for 10 mins prior to being measured. Luminescence was recorded using BioTek Synergy™ Neo2 Multimode Microplate Reader with a gain of 105. Data was plotted in GraphPad Prism 7.

***Cellular Toxicity with Adherent Cells (HEK-293T & U87)***

Adherent cells were plated in 96 well opaque plates at a density of 10,000 cells/well (100 µL per well of a 100,000 cells/mL stock). 200 µL of PBS was added to surrounding wells. Cells were allowed to adhere overnight in cell incubator. Cell media was then carefully removed from the plate. Cells were then treated with 100 µL of 1X target concentration of compound stocks prepared at 0.5% DMSO solution in cell media (1% FBS), in triplicate. Cell plate was placed into the incubator for 48 hours. After 48 hours, cell plate was removed and brought to r.t. for 30 mins. 90 µL of Cell Titer-Glo® luminescent reagent was added to each well. Plate was gently shaken for 2 minutes. Plate was then protected from light for 10 mins prior to being measured. Luminescence was recorded using BioTek Synergy™ Neo2 Multimode Microplate Reader with a gain of 105. Data was plotted in GraphPad Prism 7.

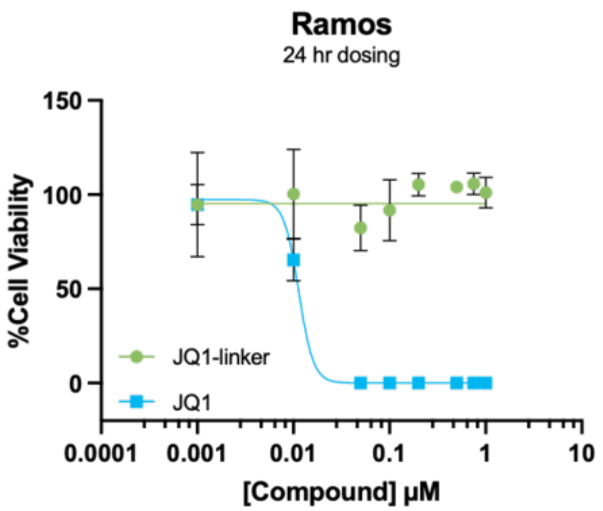


**Figure S11.** JQ1 vs. JQ1-linker control in Ramos cells for 24 hr

A

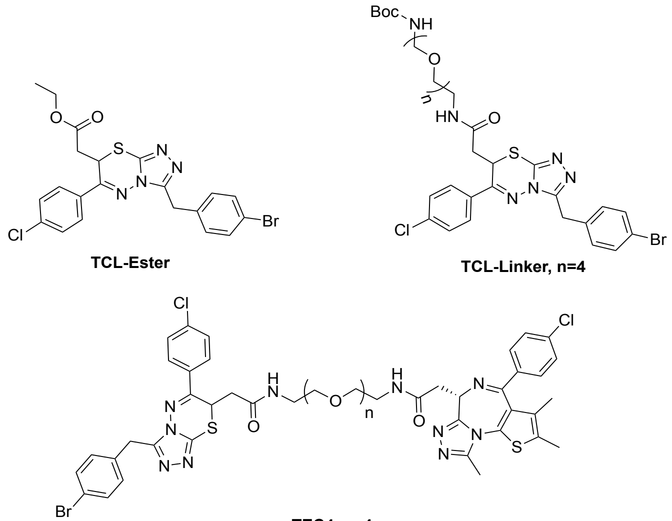


B

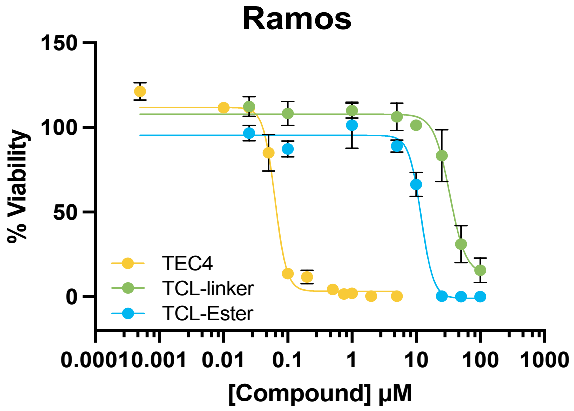


**Figure S12.** Cell Viability of TCL-Linker control (green) and TCL-ester (blue), compared to TEC4 (yellow). Addition of the linker did not make TCL more toxic indicating that incorporation of JQ1 into the bifunctional molecule to interact with BRD4 for degradation is required to be the most toxic to Ramos B-cells.

#### Appendix: $^{1}$H and $^{13}$C NMRs for TEC Degrader Series

**TEC2**

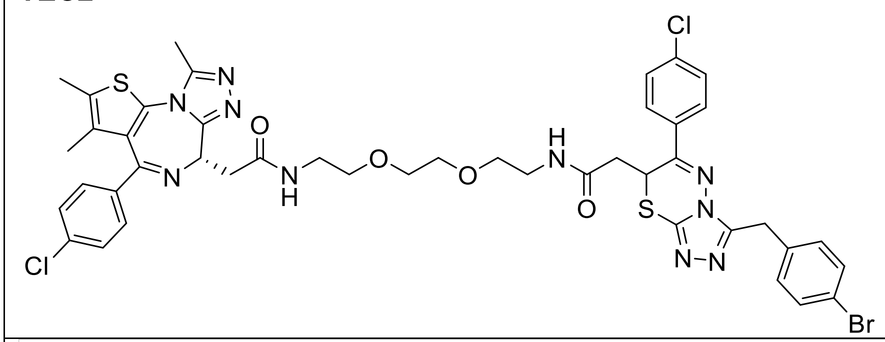

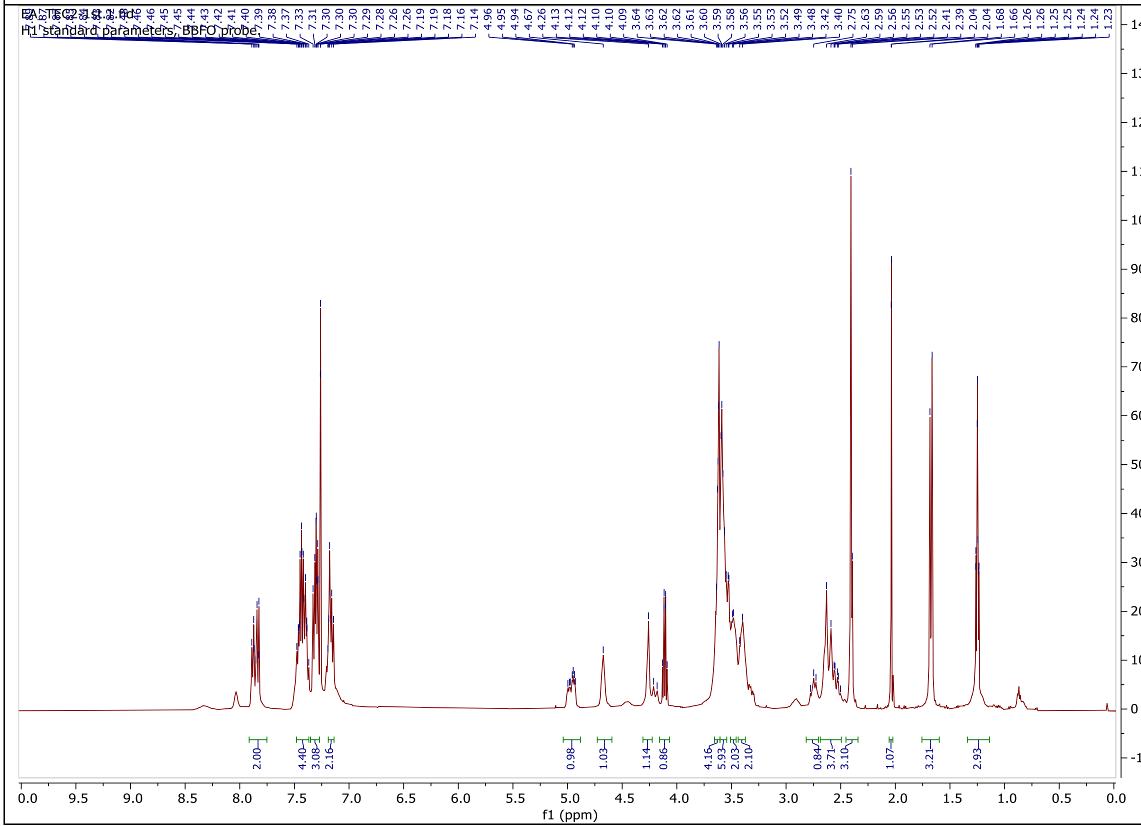

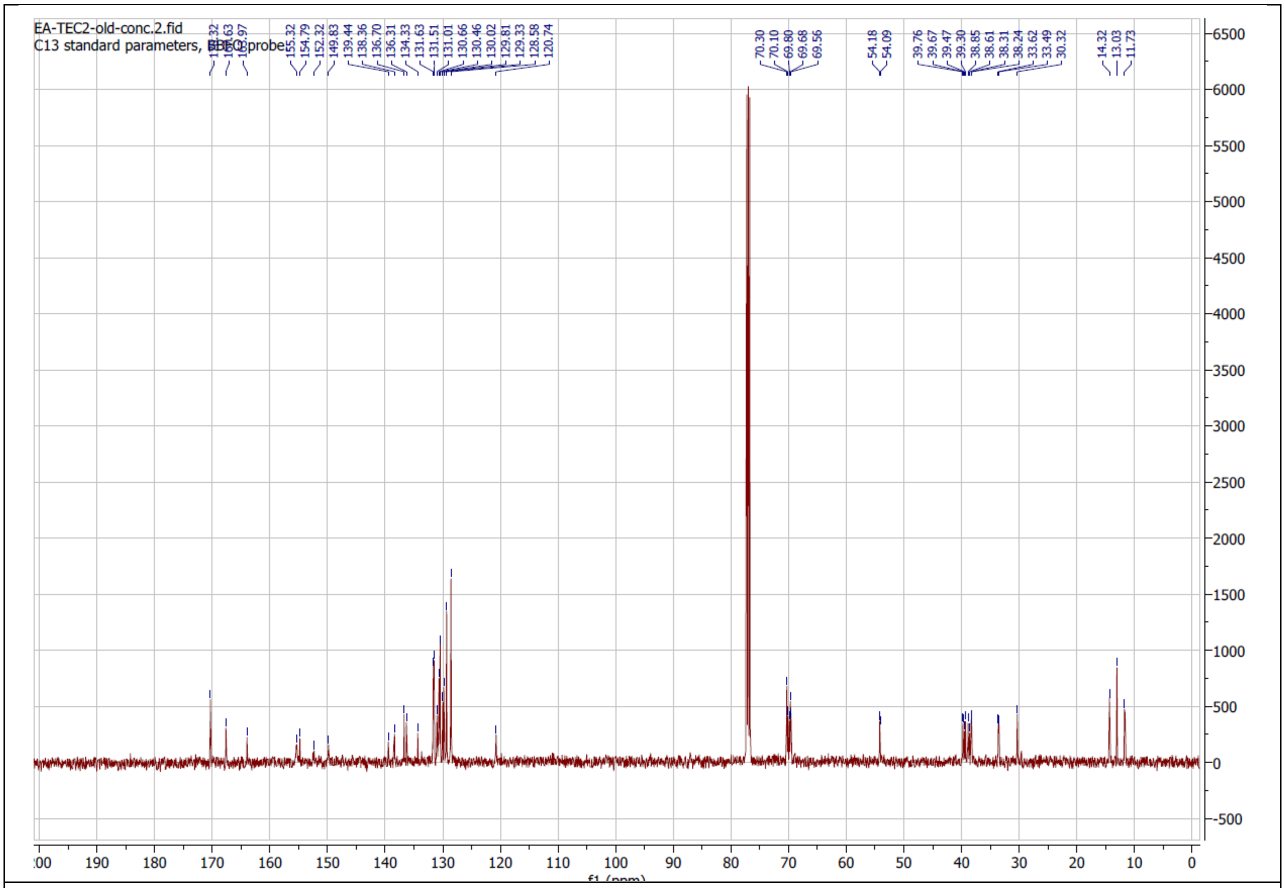

EA-TEC2-old-conc.2.fid
C13 standard parameters
155.32
154.79
152.32
149.83
139.44
138.36
136.70
136.31
134.33
131.63
131.51
131.01
130.66
130.46
130.02
129.81
129.33
128.58
120.74
70.30
70.10
69.80
69.68
69.56
54.18
54.09
39.76
39.67
39.47
39.30
38.85
38.61
38.31
38.24
33.62
33.49
30.32
14.32
13.03
11.73
6500
6000
5500
5000
4500
4000
3500
3000
2500
2000
1500
1000
500
0
-500
190
180
170
160
150
140
130
120
110
100
90
80
70
60
50
40
30
20
10
0

### TEC3

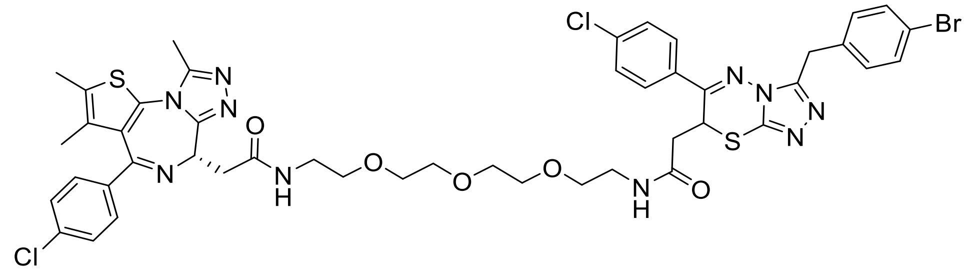

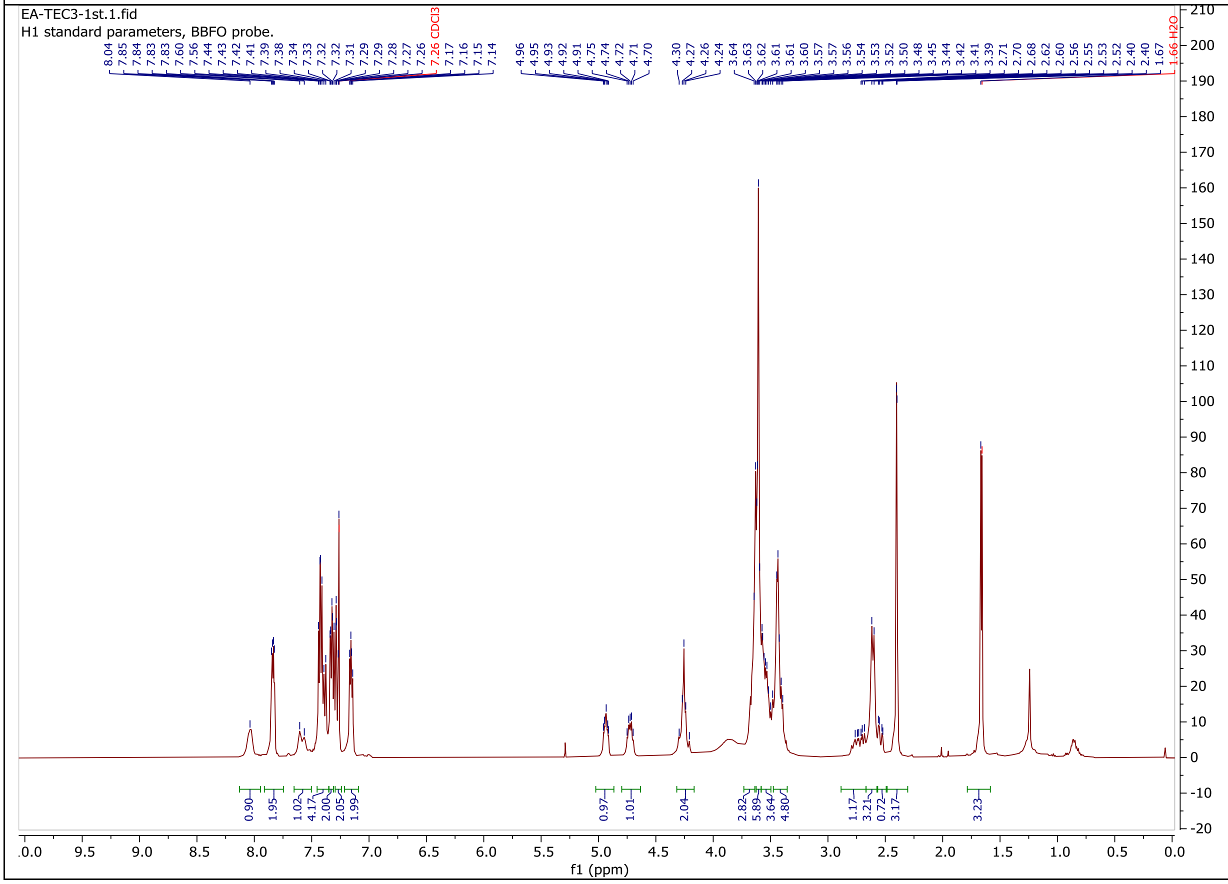
EA-TEC3-1st.1.fid
H1 standard parameters, BBFO probe.
7.26 CDCl3
1.66 H2O
f1 (ppm)

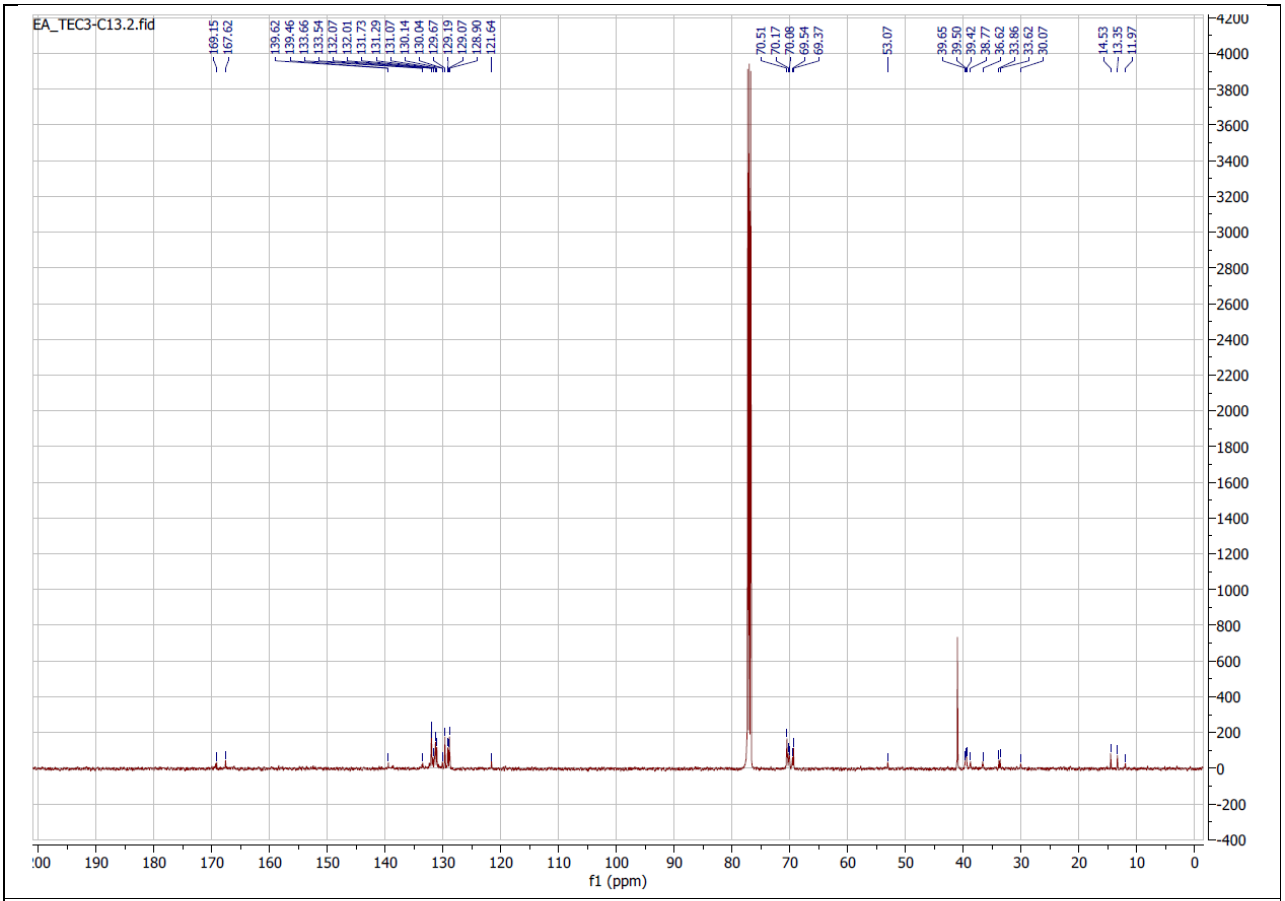
EA_TEC3-C13.2.fid
169.15
167.62
139.62
139.46
133.66
133.54
132.07
132.01
131.73
131.29
131.07
130.14
130.04
129.67
129.19
129.07
128.90
121.64
70.51
70.17
70.08
69.54
69.37
53.07
39.65
39.50
39.42
38.77
36.62
33.86
33.62
30.07
14.53
13.35
11.97
f1 (ppm)

**TEC4**

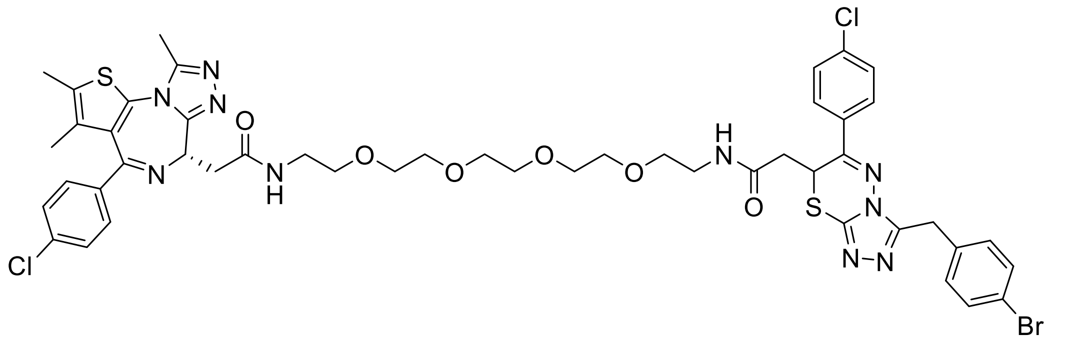

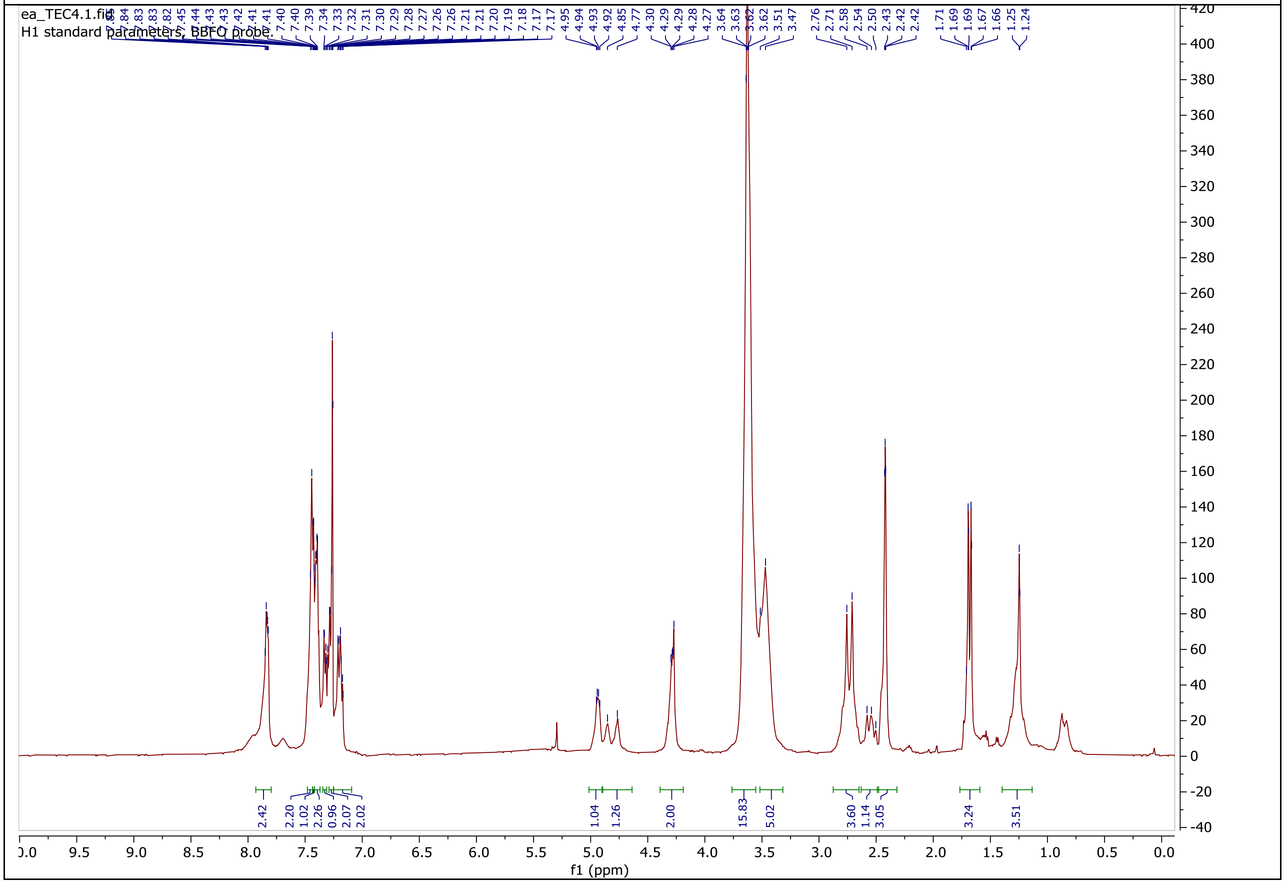

ea_TEC4.1.fid
H1 standard parameters, BBFO probe.
7.84 7.83 7.83 7.82 7.45 7.44 7.43 7.43 7.42 7.41 7.41 7.40 7.39 7.34 7.33 7.32 7.31 7.30 7.29 7.28 7.27 7.26 7.26 7.21 7.21 7.20 7.19 7.18 7.17 7.17 4.95 4.94 4.93 4.92 4.85 4.77 4.30 4.29 4.29 4.28 4.27 3.64 3.63 3.62 3.62 3.51 3.47 2.76 2.71 2.58 2.54 2.50 2.43 2.42 2.42 1.71 1.69 1.69 1.67 1.66 1.25 1.24
2.42 2.20 1.02 2.26 0.96 2.07 2.02 1.04 1.26 2.00 15.83 5.02 3.60 1.14 3.05 3.24 3.51
f1 (ppm)

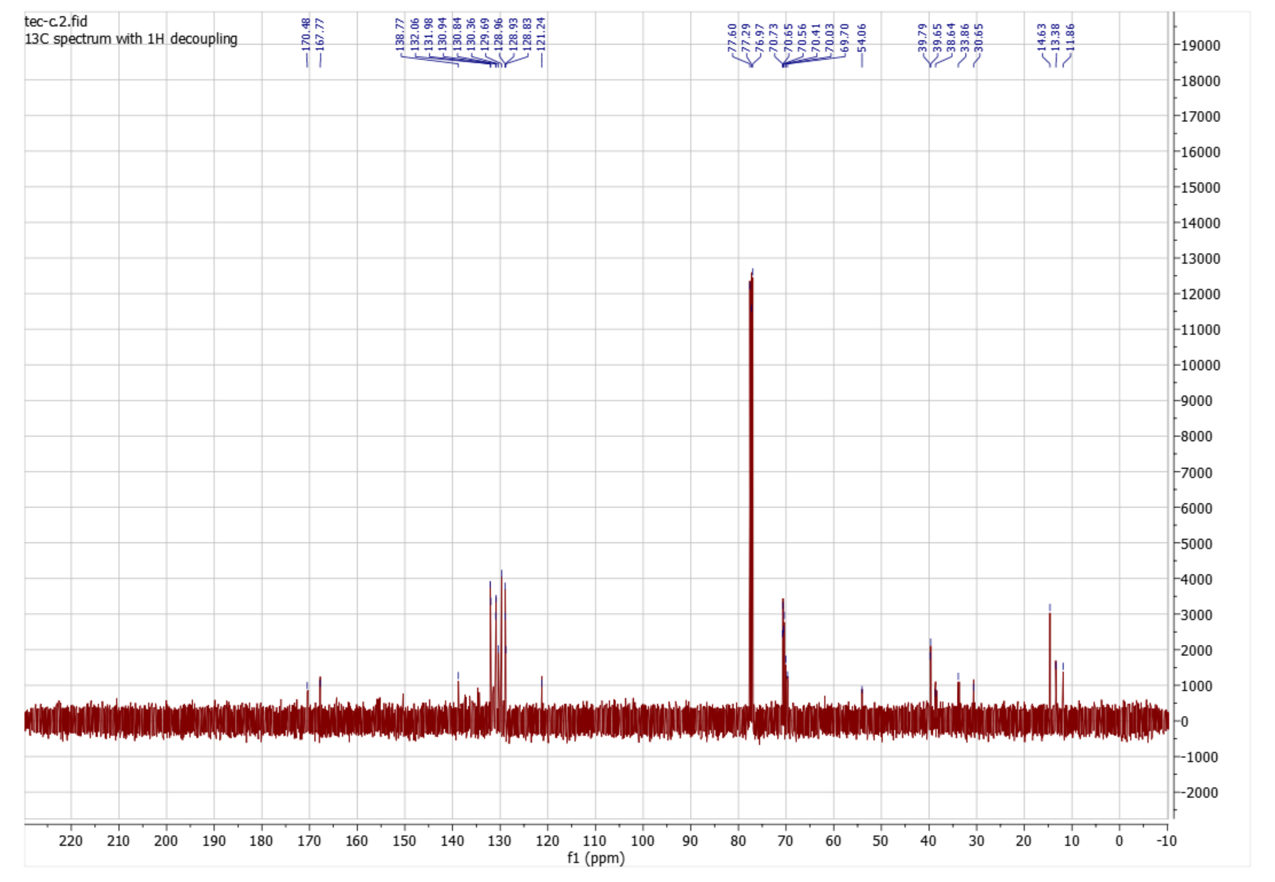
tec-c.2.fid
13C spectrum with 1H decoupling
170.48
167.77
138.77
132.06
131.98
130.94
130.84
130.36
129.69
128.96
128.93
128.83
121.24
77.60
77.29
76.97
70.73
70.65
70.56
70.41
70.03
69.70
54.06
39.79
39.65
38.64
33.86
30.65
14.63
13.38
11.86
f1 (ppm)

### TEC5

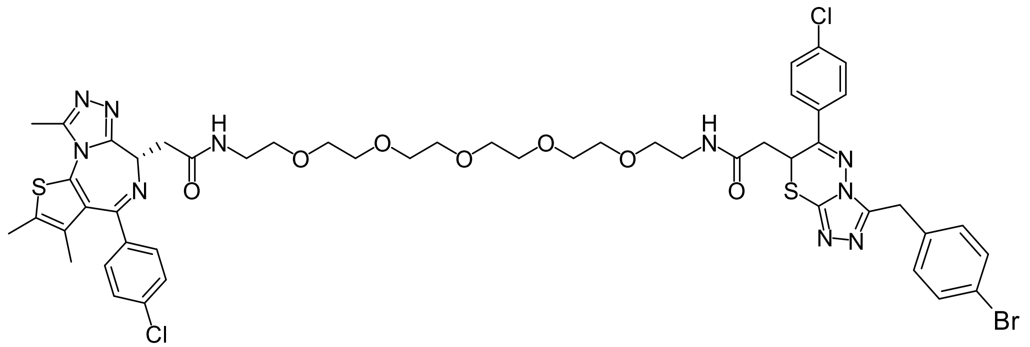

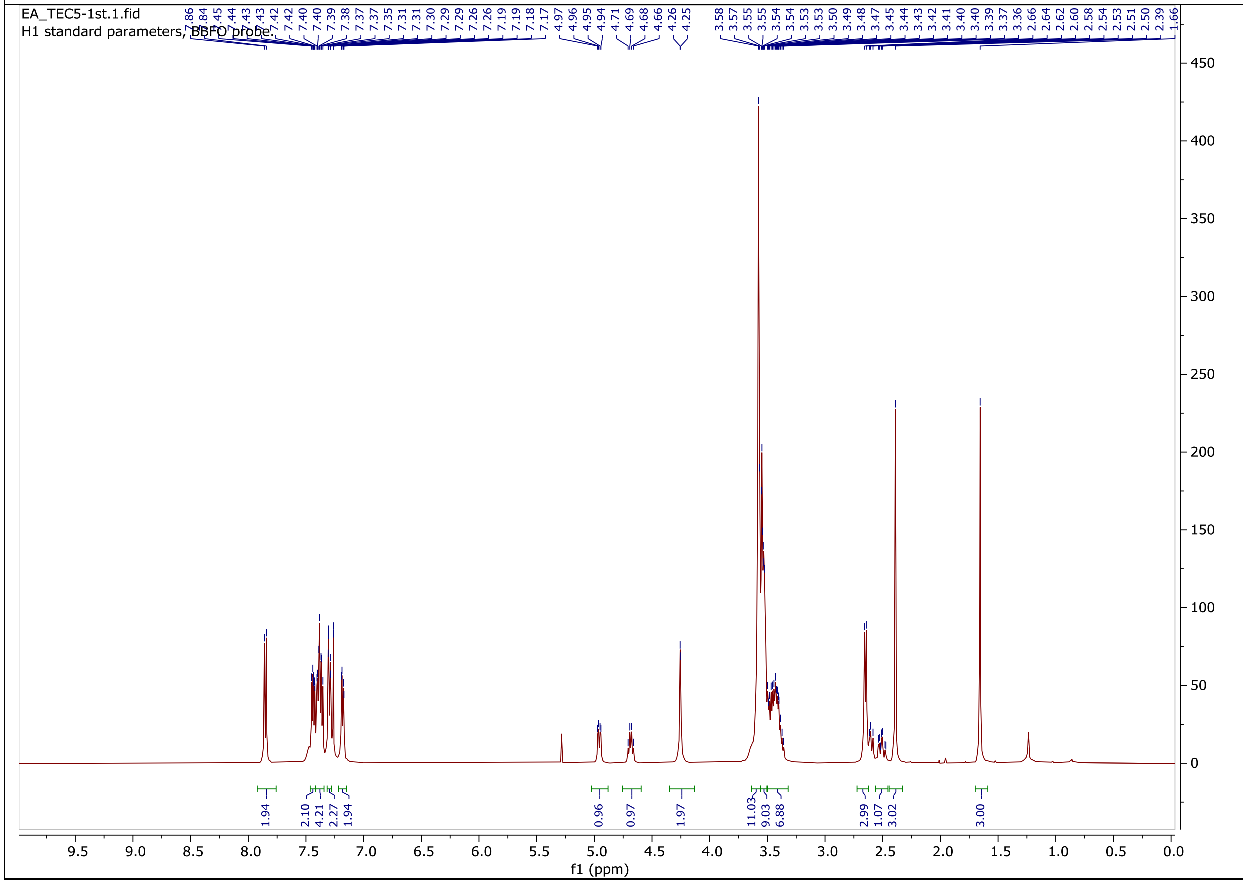
EA_TEC5-1st.1.fid
H1 standard parameters, BBFO probe
7.86
7.84
7.45
7.44
7.43
7.43
7.42
7.40
7.40
7.39
7.38
7.37
7.37
7.35
7.31
7.31
7.30
7.29
7.29
7.26
7.26
7.19
7.19
7.18
7.17
4.97
4.96
4.95
4.94
4.71
4.69
4.68
4.66
4.26
4.25
3.58
3.57
3.55
3.55
3.54
3.54
3.53
3.53
3.50
3.49
3.48
3.47
3.45
3.44
3.43
3.42
3.41
3.40
3.40
3.39
3.37
3.36
2.66
2.64
2.62
2.60
2.58
2.54
2.53
2.51
2.50
2.39
1.66
1.94
2.10
4.21
2.27
1.94
0.96
0.97
1.97
11.03
9.03
6.88
2.99
1.07
3.02
3.00
450
400
350
300
250
200
150
100
50
0
9.5
9.0
8.5
8.0
7.5
7.0
6.5
6.0
5.5
5.0
4.5
4.0
3.5
3.0
2.5
2.0
1.5
1.0
0.5
0.0
f1 (ppm)

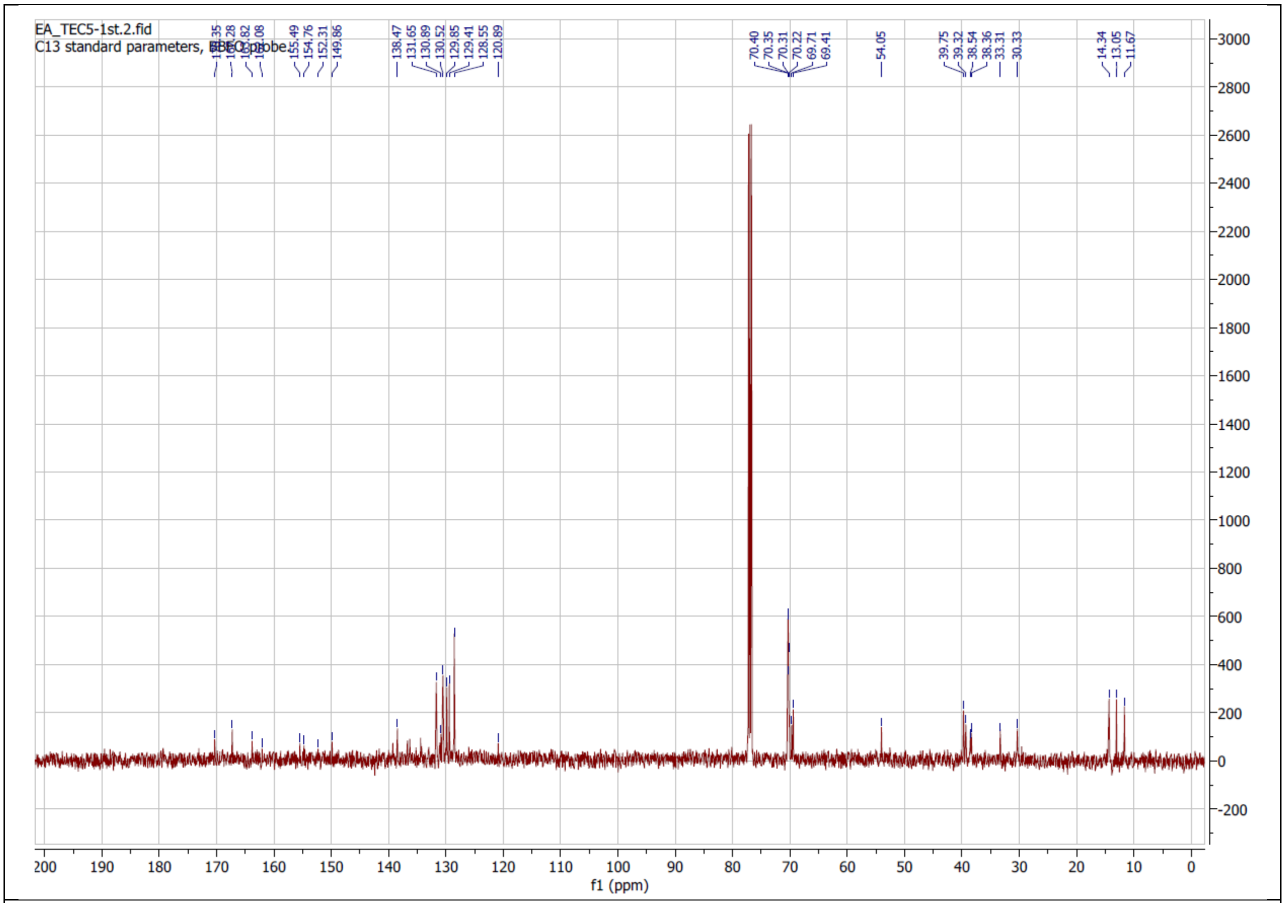

EA_TEC5-1st.2.fid
155.49
154.76
152.31
149.86
138.47
131.65
130.89
130.52
129.85
129.41
128.55
120.89
70.40
70.35
70.31
70.22
69.71
69.41
54.05
39.75
39.32
38.54
38.36
33.31
30.33
14.34
13.05
11.67
f1 (ppm)

**TEC6**

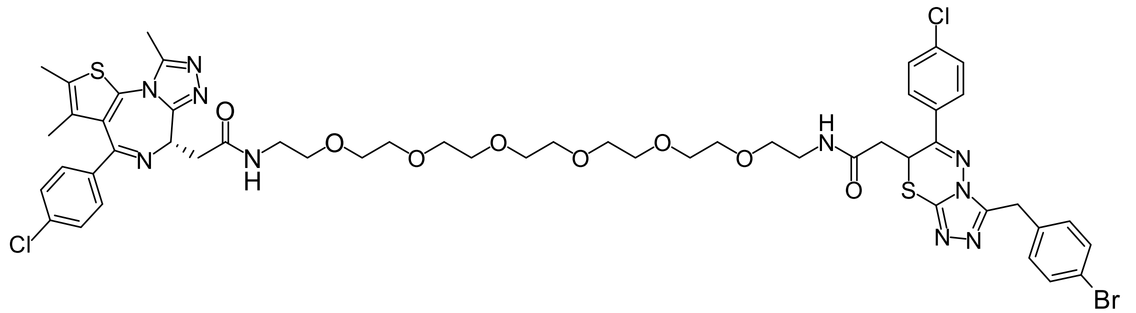
S
N
N
N
O
N
H
O
O
O
O
O
O
H
N
O
Cl
N
N
S
N
N
N
Cl
Br

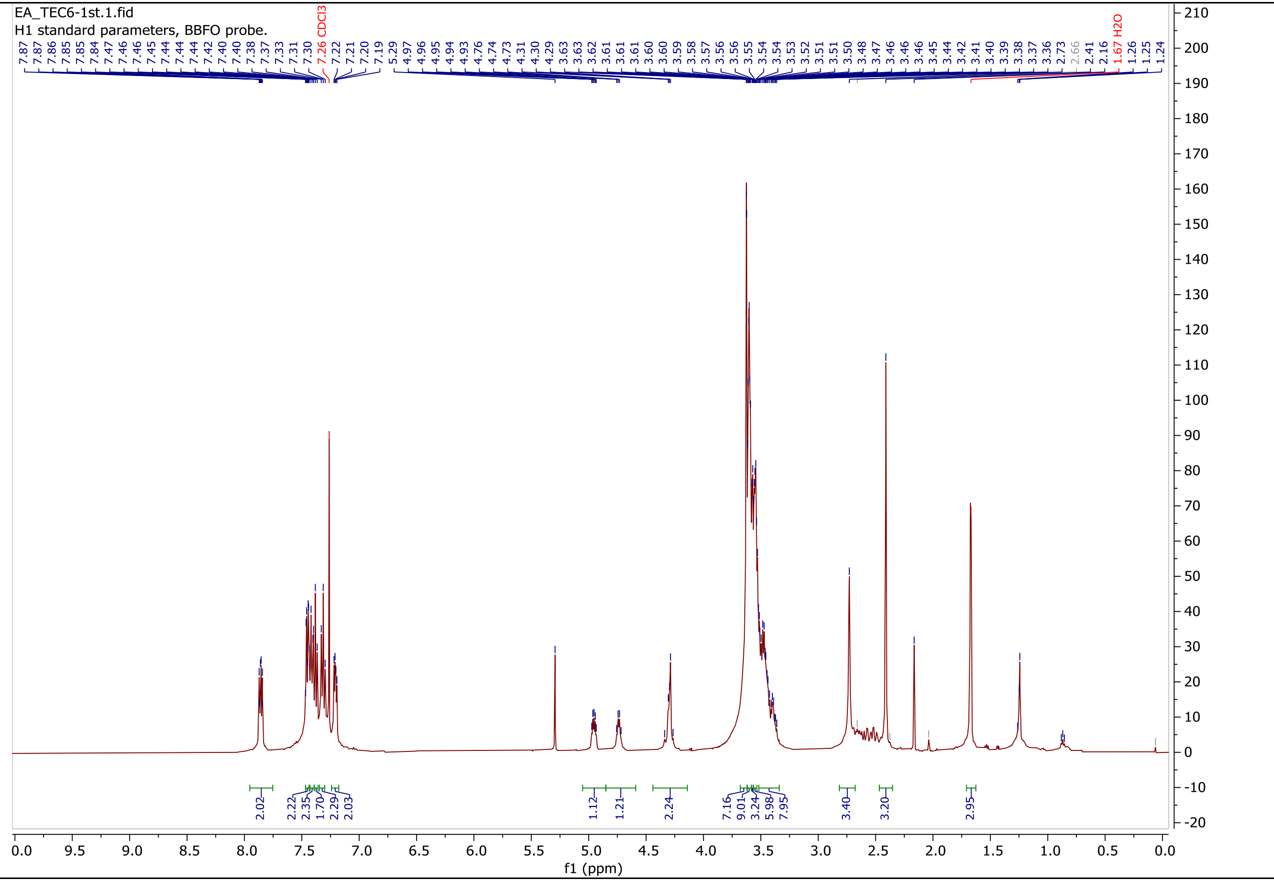
EA_TEC6-1st.1.fid
H1 standard parameters, BBFO probe.
7.87 7.87 7.86 7.85 7.85 7.84 7.47 7.46 7.46 7.45 7.44 7.44 7.44 7.42 7.40 7.40 7.38 7.37 7.33 7.31 7.30 7.26 CDCl3 7.22 7.21 7.20 7.19 5.29 4.97 4.96 4.95 4.94 4.93 4.76 4.74 4.73 4.31 4.30 4.29 3.63 3.63 3.62 3.61 3.61 3.61 3.60 3.59 3.58 3.57 3.56 3.56 3.55 3.55 3.54 3.54 3.53 3.52 3.51 3.51 3.50 3.48 3.47 3.46 3.46 3.46 3.45 3.44 3.42 3.41 3.40 3.39 3.38 3.37 3.36 2.73 2.66 2.41 2.16 1.67 H2O 1.26 1.25 1.24
2.02 2.22 2.35 1.70 2.29 2.03 1.12 1.21 2.24 7.16 9.01 3.24 5.98 7.95 3.40 3.20 2.95
f1 (ppm)

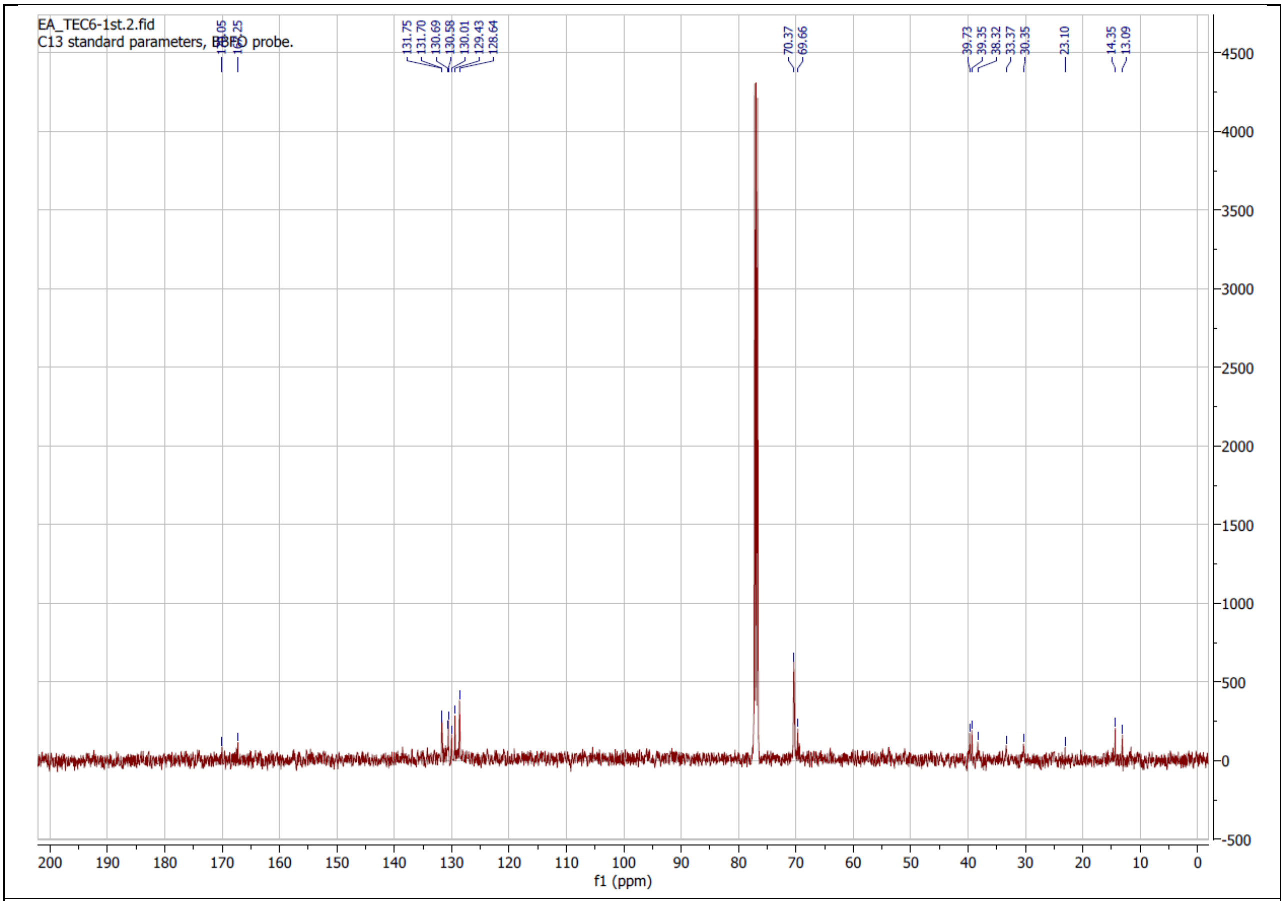
EA_TEC6-1st.2.fid
C13 standard parameters, BBFO probe.
170.05
167.25
131.75
131.70
130.69
130.58
130.01
129.43
128.64
70.37
69.66
39.73
39.35
38.32
33.37
30.35
23.10
14.35
13.09
f1 (ppm)

**TEC7**

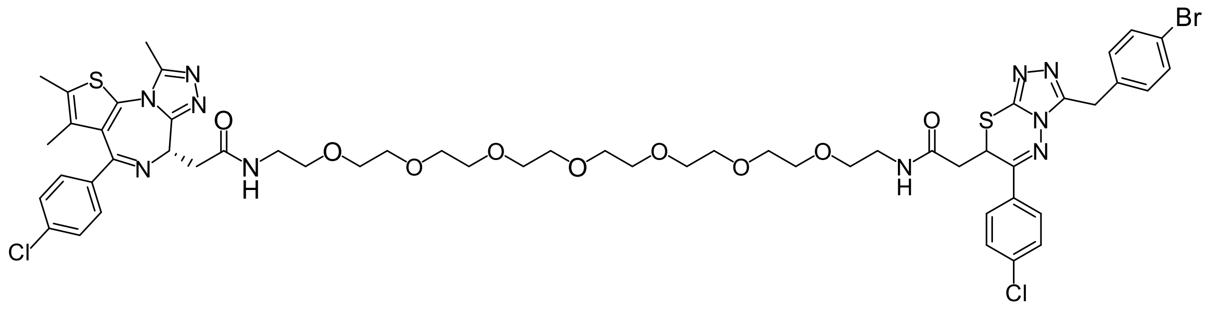

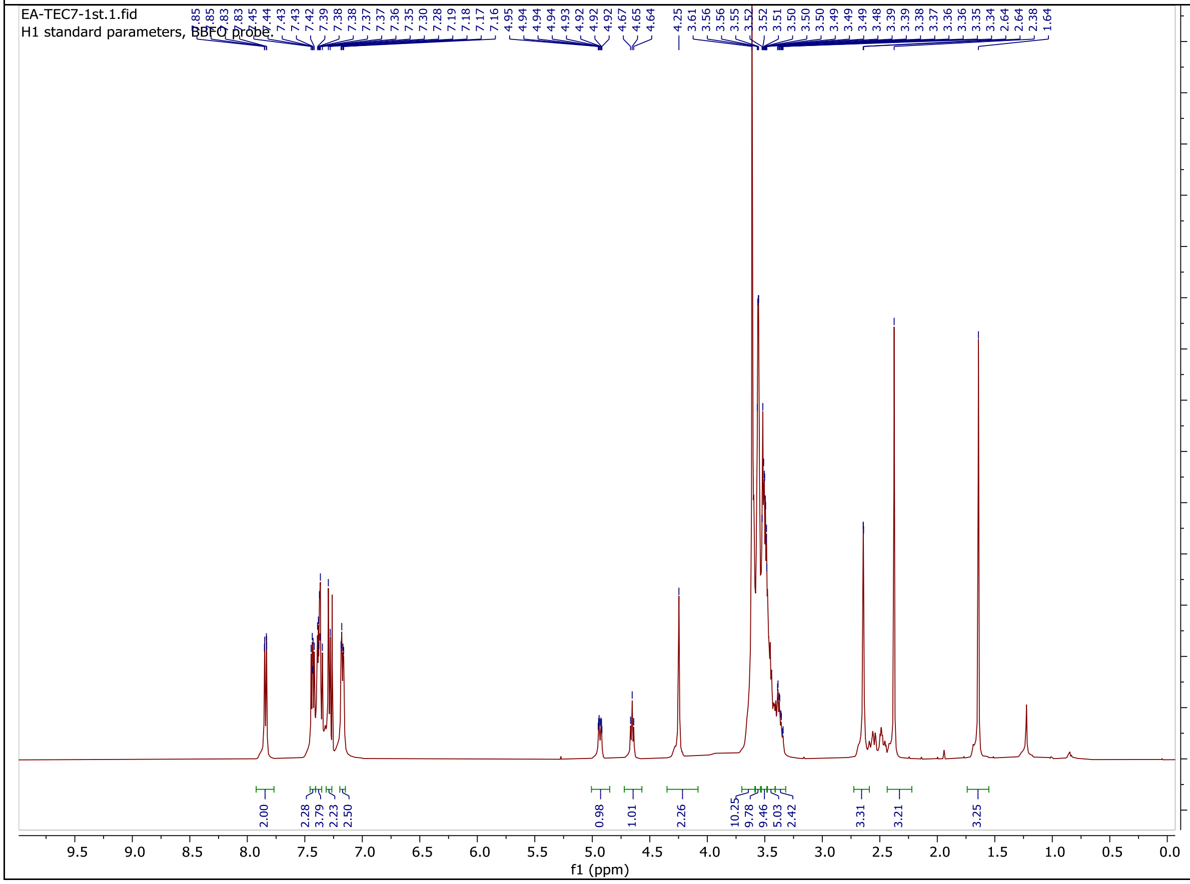

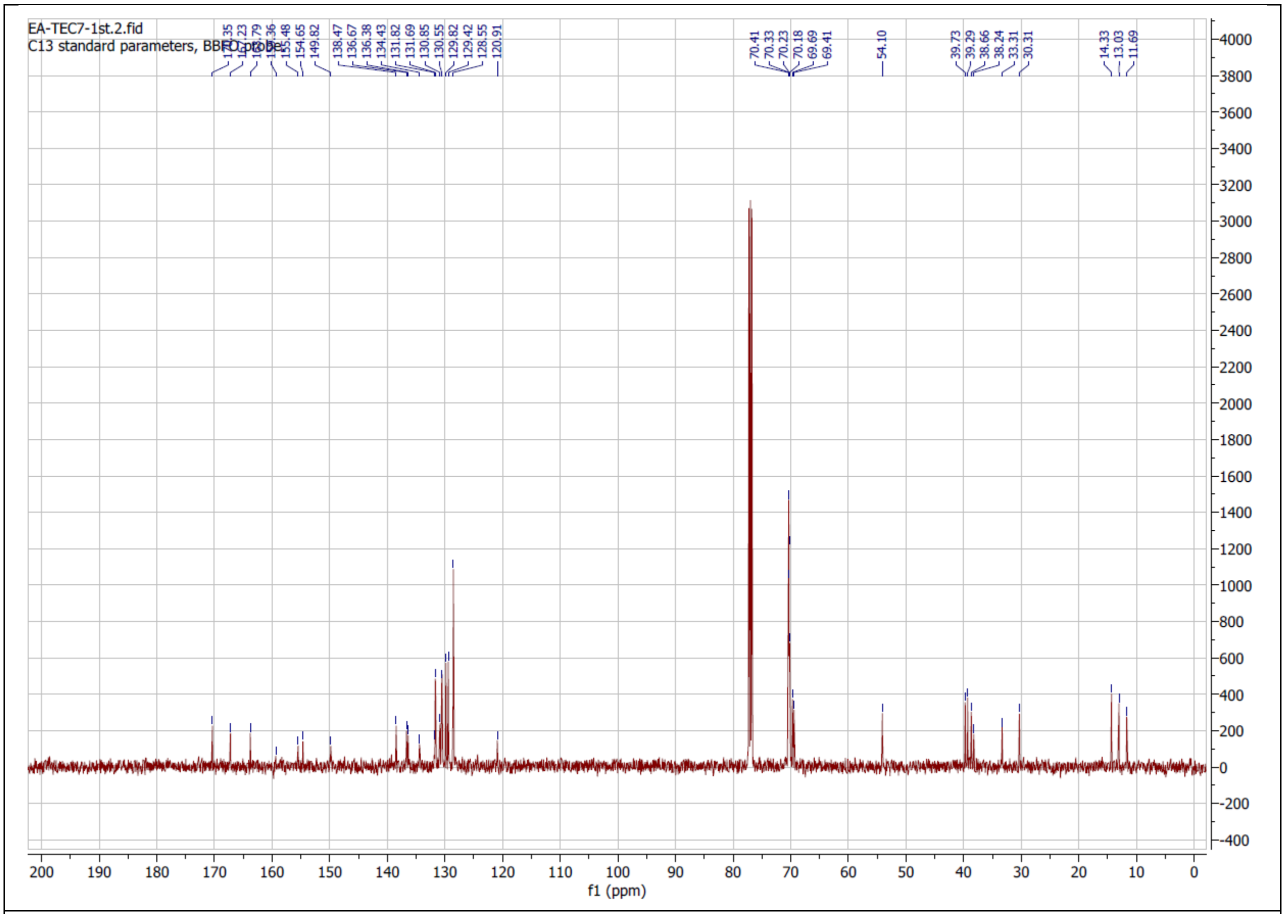
EA-TEC7-1st.2.fid
C13 standard parameters, BBFO probe
155.48
154.65
149.82
138.47
136.67
136.38
134.43
131.82
131.69
130.85
130.55
129.82
129.42
128.55
120.91
70.41
70.33
70.23
70.18
69.69
69.41
54.10
39.73
39.29
38.66
38.24
33.31
30.31
14.33
13.03
11.69
f1 (ppm)

### TEC8

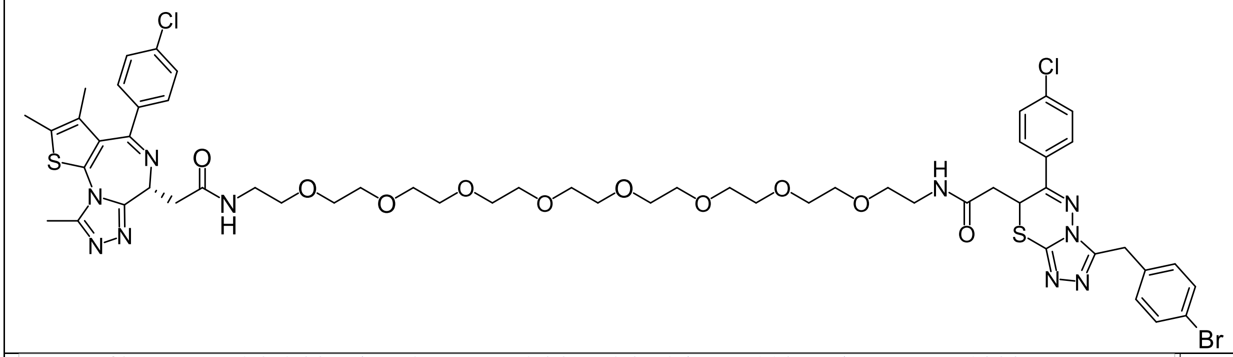

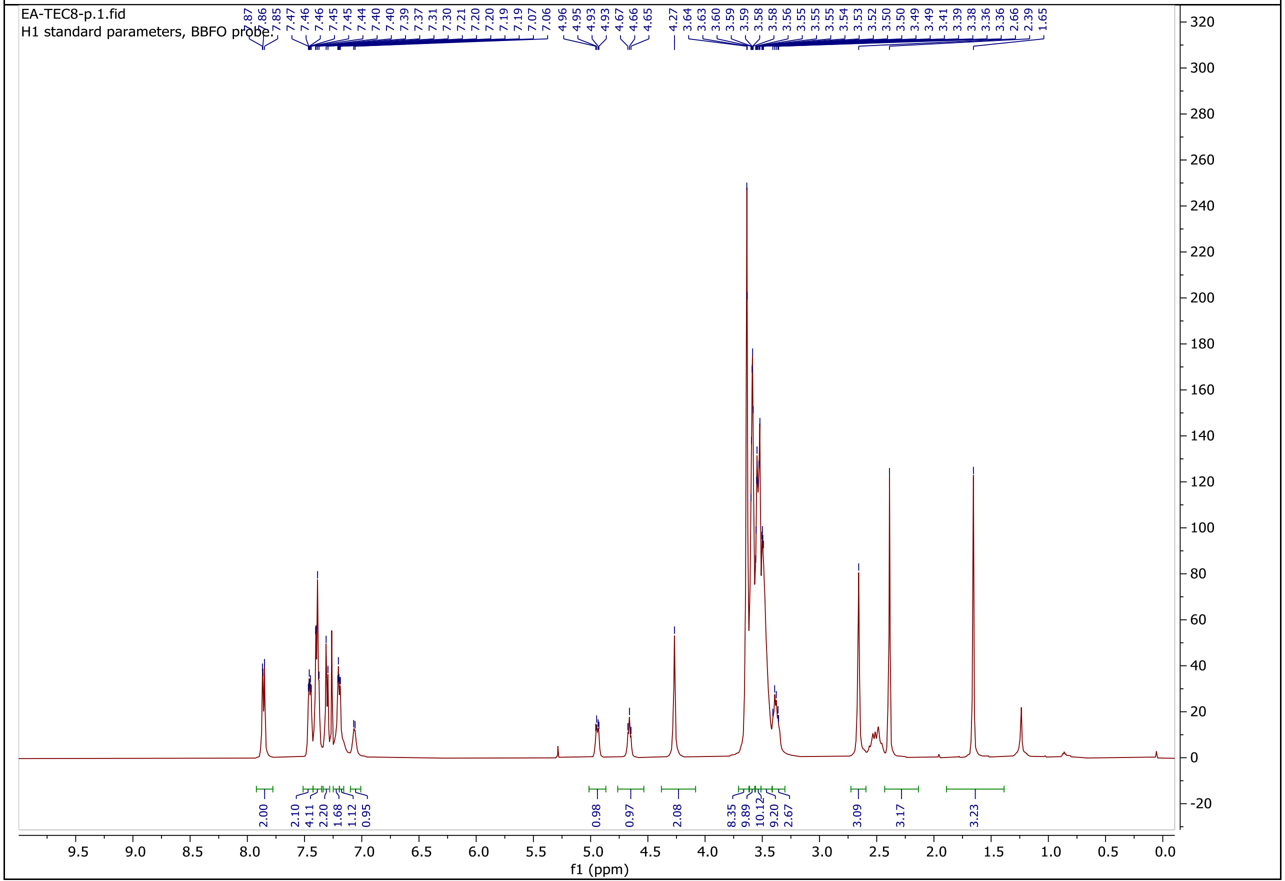

EA-TEC8-p.1.fid
H1 standard parameters, BBFO probe
7.87 7.86 7.85 7.47 7.46 7.46 7.45 7.45 7.44 7.40 7.39 7.37 7.31 7.30 7.21 7.20 7.20 7.19 7.19 7.07 7.06 4.96 4.95 4.93 4.93 4.67 4.66 4.65 4.27 3.64 3.63 3.60 3.59 3.59 3.58 3.58 3.56 3.55 3.55 3.54 3.53 3.52 3.50 3.50 3.49 3.49 3.41 3.39 3.38 3.36 3.36 2.66 2.39 1.65
2.00 2.10 4.11 2.20 1.68 1.12 0.95 0.98 0.97 2.08 8.35 9.89 10.12 9.20 2.67 3.09 3.17 3.23
f1 (ppm)

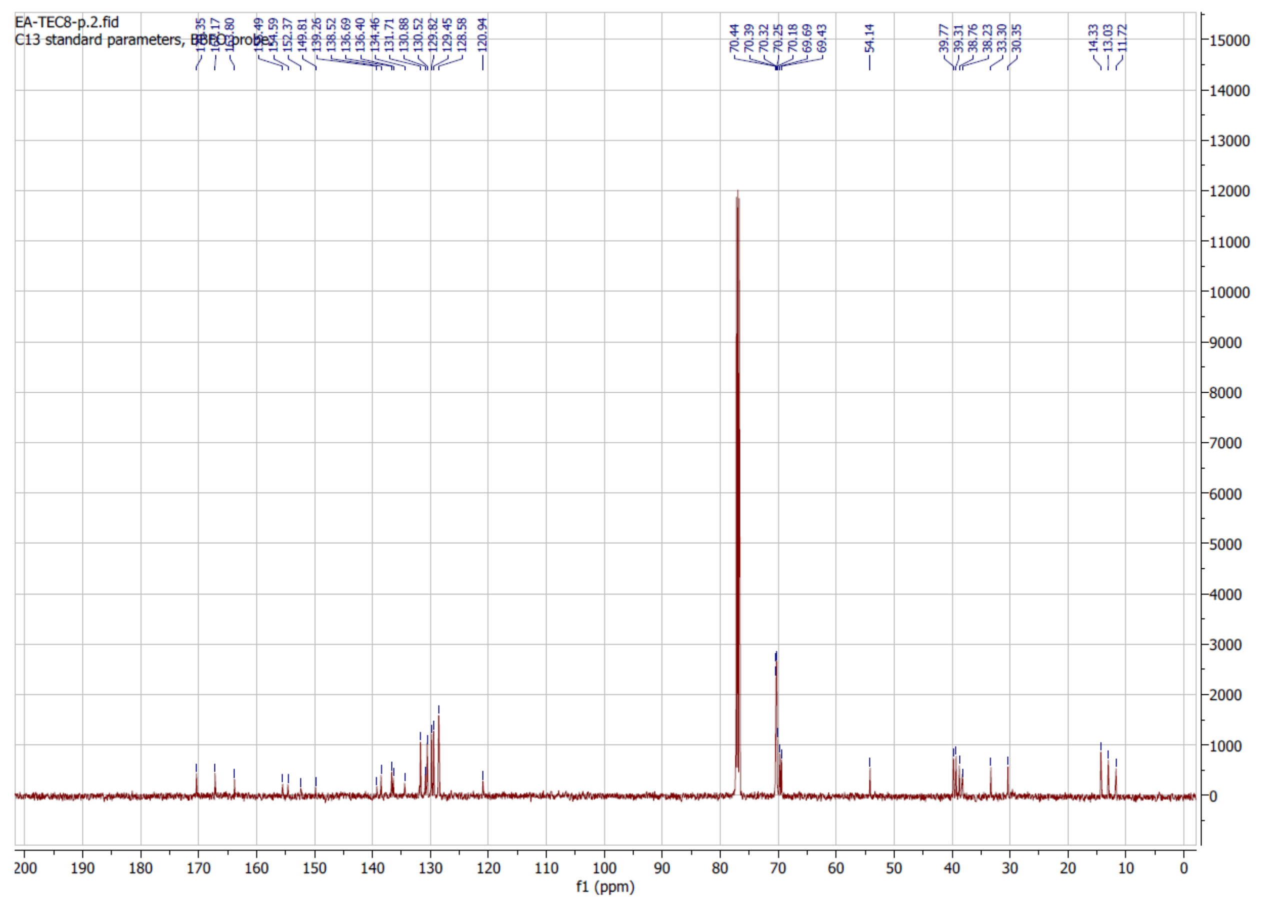

EA-TEC8-p.2.fid
C13 standard parameters, probe
152.37
149.81
139.26
138.52
136.69
136.40
134.46
131.71
130.88
130.52
129.82
129.45
128.58
120.94
70.44
70.39
70.32
70.25
70.18
69.69
69.43
54.14
39.77
39.31
38.76
38.23
33.30
30.35
14.33
13.03
11.72
f1 (ppm)

### TEC9

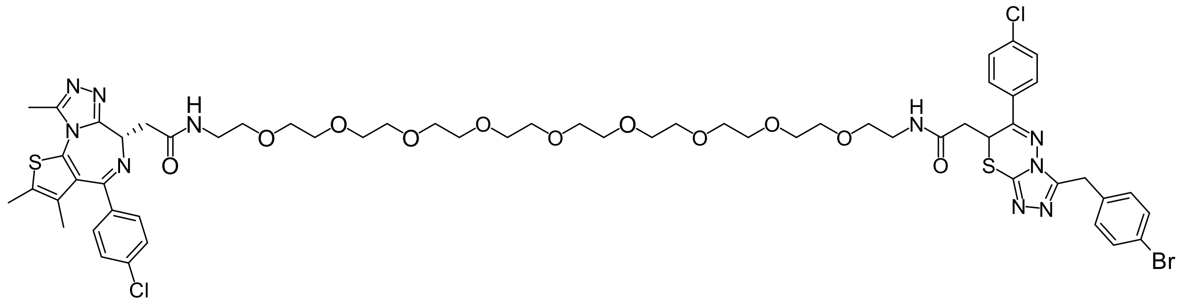

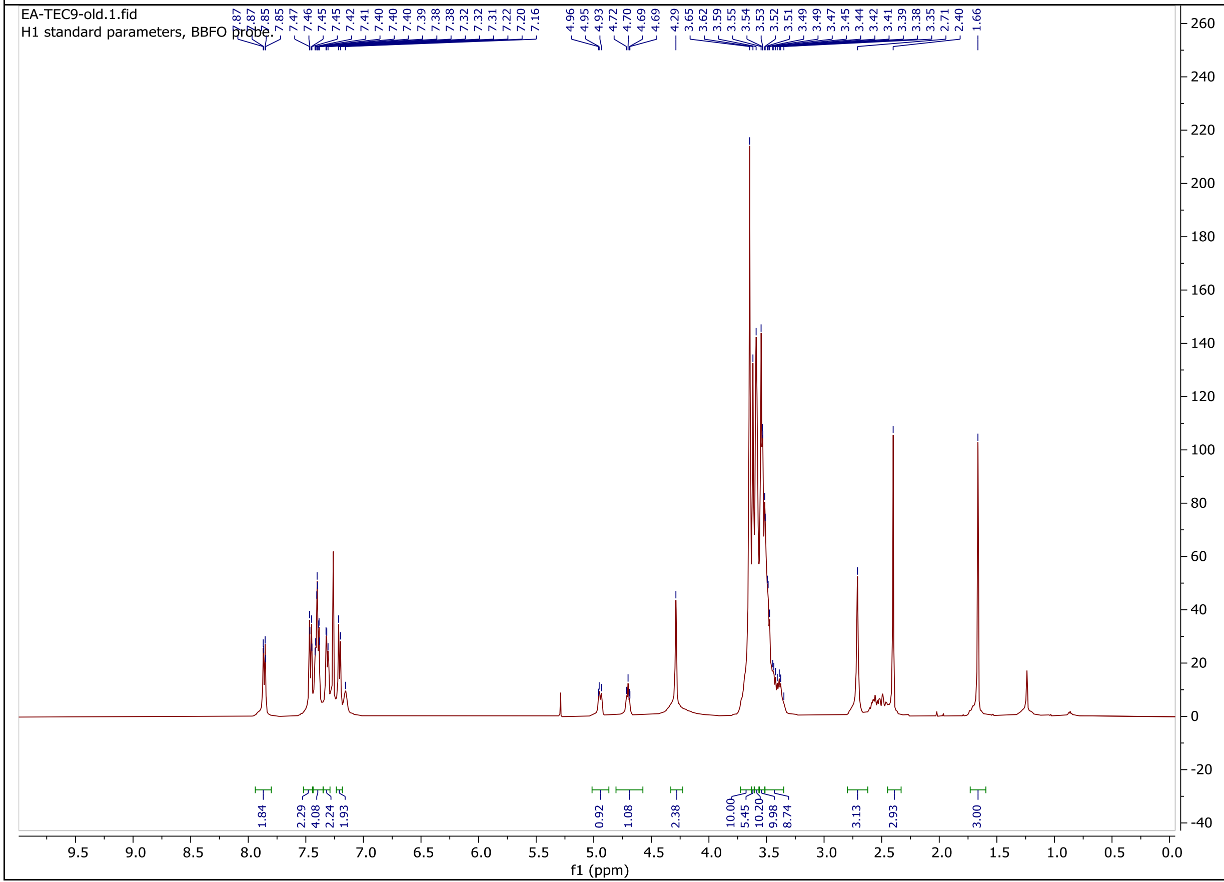
EA-TEC9-old.1.fid
H1 standard parameters, BBFO probe.
7.87 7.87 7.85 7.85 7.47 7.46 7.45 7.45 7.42 7.41 7.40 7.40 7.40 7.39 7.38 7.38 7.32 7.32 7.31 7.22 7.20 7.16
4.96 4.95 4.93 4.72 4.70 4.69 4.69 4.29
3.65 3.62 3.59 3.55 3.54 3.53 3.52 3.51 3.49 3.49 3.47 3.45 3.44 3.42 3.41 3.39 3.38 3.35 2.71 2.40 1.66
1.84 2.29 4.08 2.24 1.93 0.92 1.08 2.38 10.00 5.45 10.20 9.98 8.74 3.13 2.93 3.00
f1 (ppm)

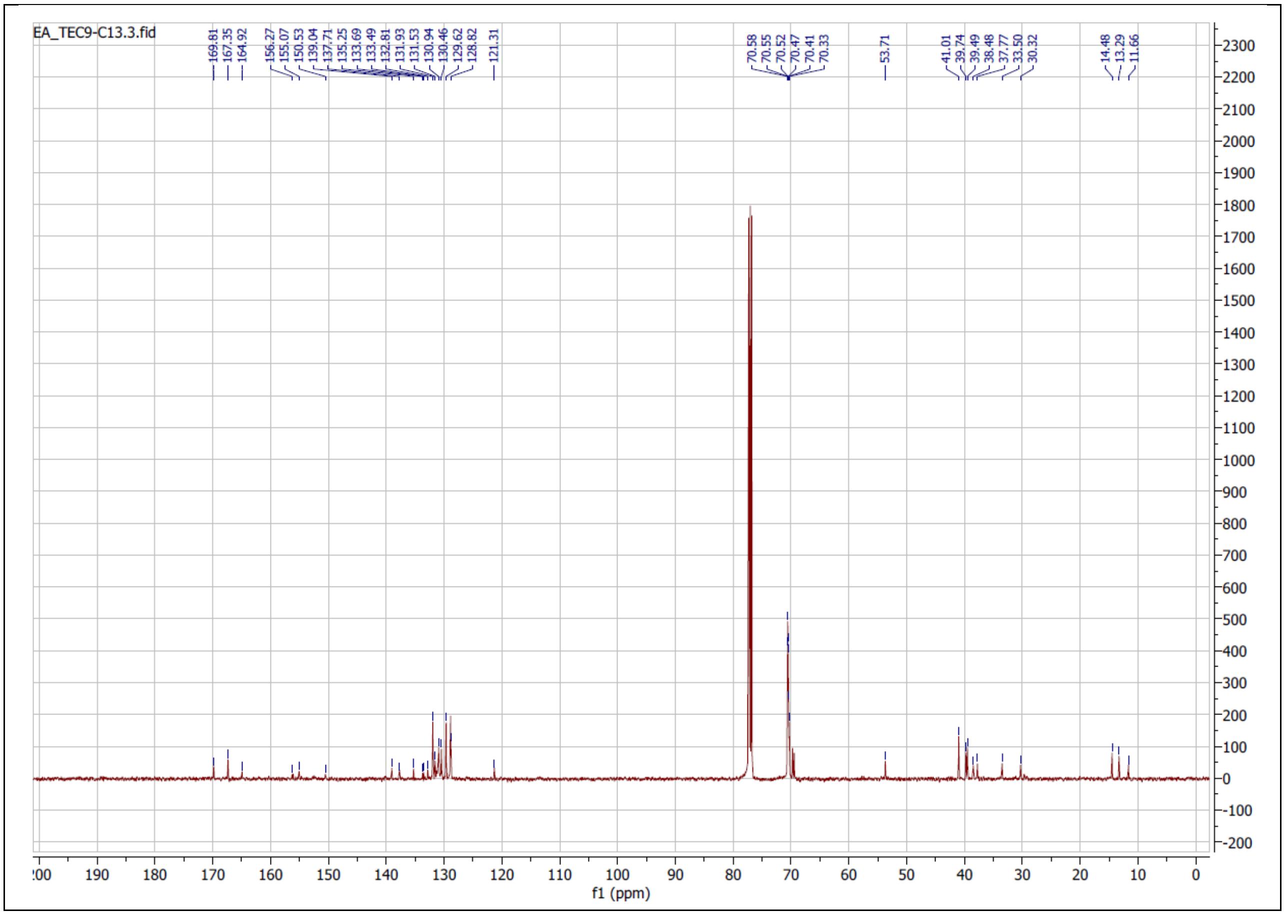
EA_TEC9-C13.3.fid
169.81
167.35
164.92
156.27
155.07
150.53
139.04
137.71
135.25
133.69
133.49
132.81
131.93
131.53
130.94
130.46
129.62
128.82
121.31
70.58
70.55
70.52
70.47
70.41
70.33
53.71
41.01
39.74
39.49
38.48
37.77
33.50
30.32
14.48
13.29
11.66
f1 (ppm)
